# A Bayesian implementation of the multispecies coalescent model with introgression for comparative genomic analysis

**DOI:** 10.1101/766741

**Authors:** Thomas Flouris, Xiyun Jiao, Bruce Rannala, Ziheng Yang

## Abstract

Recent analyses suggest that cross-species gene flow or introgression is common in nature, especially during species divergences. Genomic sequence data can be used to infer introgression events and to estimate the timing and intensity of introgression, providing an important means to advance our understanding of the role of gene flow in speciation. Here we implement the multispecies-coalescent-with-introgression (MSci) model, an extension of the multispecies-coalescent (MSC) model to incorporate introgression, in our Bayesian Markov chain Monte Carlo (MCMC) program BPP. The MSci model accommodates deep coalescence (or incomplete lineage sorting) and introgression and provides a natural framework for inference using genomic sequence data. Computer simulation confirms the good statistical properties of the method, although hundreds or thousands of loci are typically needed to estimate introgression probabilities reliably. Re-analysis of datasets from the purple cone spruce confirms the hypothesis of homoploid hybrid speciation. We estimated the introgression probability using the genomic sequence data from six mosquito species in the *Anopheles gambiae* species complex, which varies considerably across the genome, likely driven by differential selection against introgressed alleles.

## Introduction

A number of recent studies have revealed cross-species hybridization/introgression in a variety of species ranging from Arabidopsis (Arnold *et al*., 2016), butterflies (Martin *et al*., 2013), corals (Mao *et al*., 2018), and birds (Ellegren *et al*., 2012) to mammals such as bears (Kumar *et al*., 2017; Liu *et al*., 2014), cattle (Wu *et al*., 2018), gibbons (Chan *et al*., 2013; Shi and Yang, 2018), and hominins (Nielsen *et al*., 2017). Introgression may play an important role in speciation (Harrison and Larson, 2014; Mallet *et al*., 2016; Martin and Jiggins, 2017). Inference of introgression and estimation of migration rates can contribute to our understanding of the speciation process (Mallet *et al*., 2016; Martin and Jiggins, 2017). Furthermore, introgression and deep coalescence (or incomplete lineage sorting, ILS) are two major challenges for species tree reconstruction (Fontaine *et al*., 2015; Liu *et al*., 2014; Martin *et al*., 2013).

There is a large body of literature on the use of networks to model non-treelike evolution (Huson *et al*., 2011) and several methods have been developed to detect cross-species gene flow. Most use summaries of the multi-locus sequence data such as the frequencies of estimated gene tree topologies (Solis-Lemus and Ane, 2016; Solis-Lemus *et al*., 2017) or the counts of parsimony-informative site patterns (Blischak *et al*., 2018; Durand *et al*., 2011; Green *et al*., 2010). See Degnan (2018) and Folk *et al*. (2018) for recent reviews. We focus on coalescent-based full-likelihood models for closely related species. These come in two forms. The isolation-with-migration (IM) model assumes continuous migration, with species exchanging migrants at certain rates every generation (Hey, 2010; Hey and Nielsen, 2004), while the MSci model assumes episodic introgression/hybridization (Yu *et al*., 2014). While the probability density of the gene trees under the IM (Hey, 2010) and MSci (Yu *et al*., 2014) models is straightforward to compute, developing a Bayesian MCMC program that is feasible for use with genome-scale datasets has been challenging. The space of unknown genealogical histories (including the migration/introgression histories) is large, and constraints between the species tree and the gene trees make it difficult to traverse the parameter space in the posterior. Current MCMC implementations include IMa3 (Hey, 2010; Hey *et al*., 2018) for the IM model, and *BEAST (Jones, 2019; Zhang *et al*., 2018) and PHYLONET (Wen and Nakhleh, 2018) for the MSci model. It does not appear computationally feasible to apply those programs to realistically sized datasets, with more than 200 loci, say. Moreover, current implementations (Wen and Nakhleh, 2018; Zhang *et al*., 2018) assume that the parental species become extinct when a hybrid species is formed (i.e., model A in Fig. 1), and exclude important biological scenarios of hybridization (models B-D in Fig. 1).

**FIG. 1.**
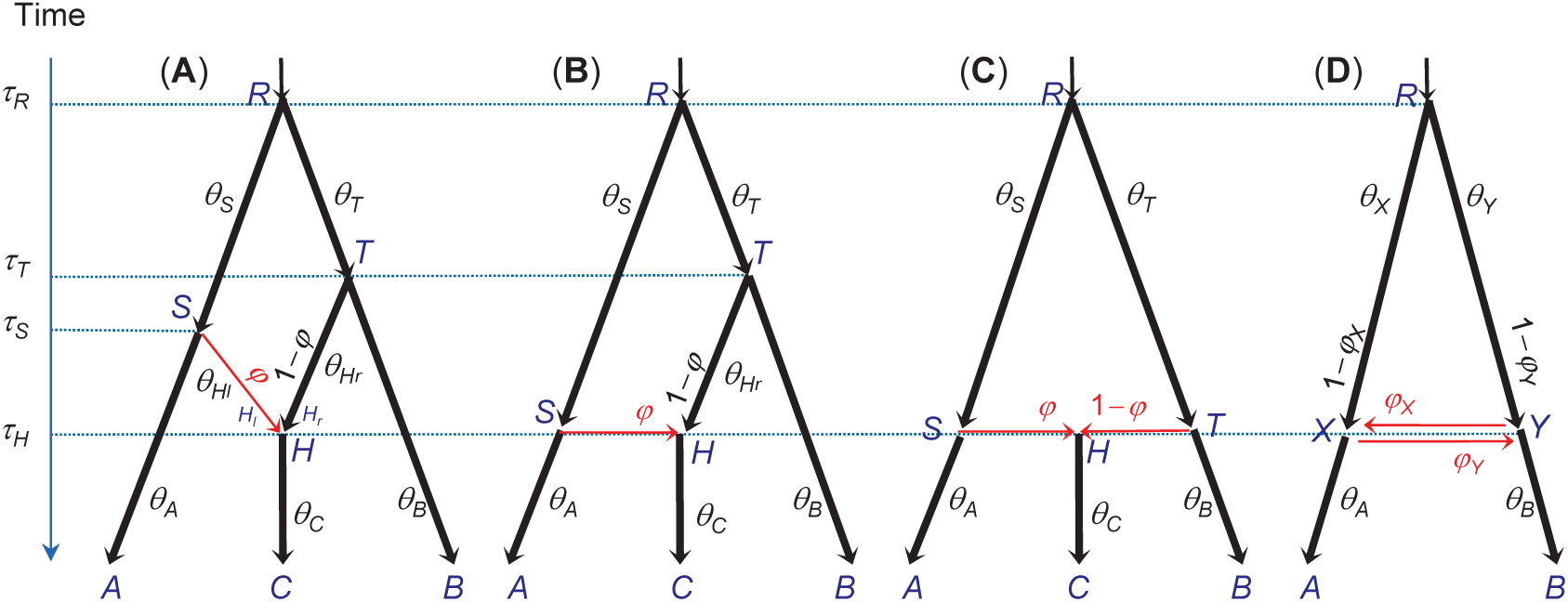
The MSci model with four different types of hybridization (introgression or admixture) events. In (A), two parental species *SH* and *TH* merge to form a hybrid species *H*, at time *τH*, leading to extinction of the parental species. In (B), there is introgression from species *RSA* to species *THC* at time *τH* = *τS*, with introgression probability *φ*. In (C), species *RSA* and *RTB* come into contact to form hybrid species *H* at time *τS* = *τH* = *τT*, which evolves into species *C*, while the two parent species become *A* and *B*. In (D), bidirectional introgression occurs between species *RXA* and *RY B* at time *τX* = *τY*, with introgression probabilities *φX* and *φY*. Parameters in the model include speciation/hybridization times (*τ*s), population sizes (*θ* s), and introgression probabilities (*φ*s). The models are represented using the extended Newick notation (Cardona *et al*., 2008), as (A)-(C): ((*A*, (*C*)*H*)*S*, (*H, B*)*T*) *R* and (D): ((*A*, (*B*)*Y*) *X*, (*X*) *Y*) *R*. Arrows indicate the direction of time from parent to child or from source to target populations.

In this paper we extend the MSC model in the BPP program (Burgess and Yang, 2008; Rannala and Yang, 2003; Yang, 2015) to accommodate introgression, resulting in the multispecies-coalescent-with-introgression model, or MSci (Degnan, 2018). The MSci model can be used to estimate species divergence times and the number, timings and intensities of introgression events. By accommodating gene flow and providing more reliable estimates of evolutionary parameters, the model may also be used in heuristic species delimitation (Jackson *et al*., 2017; Leaché *et al*., 2019). We conduct simulation to examine the statistical properties of the method, in comparison with two summary methods, SNAQ (Solis-Lemus and Ane, 2016; Solis-Lemus *et al*., 2017) and HYDE (Blischak *et al*., 2018). We apply the new method to datasets of purple cone spruce (Sun *et al*., 2014; Zhang *et al*., 2018) and genomic datasets of *Anopheles* mosquitoes (Fontaine *et al*., 2015; Thawornwattana *et al*., 2018a), to estimate the introgression probability and study its variation across the genome.

## Results

### The MSci model

We extend the MSC model (Rannala and Yang, 2003) to accommodate cross-species hybridization (or introgression) by introducing hybridization (or *H*) nodes (Fig. 1). Each *H* node has two parents (*H*_*l*_ and *H*_*r*_, for left and right) and one daughter, although the *H* node and its parents may have the same age when there is an admixture or horizontal gene transfer (Fig. 1B-C). In model A both parental species become extinct after hybridization, while model B represents an introgression from species *RSA* into *THC*. Model C represents hybrid speciation, while model D bidirectional introgression (Kubatko, 2009).

When we trace a lineage backwards in time and reach an *H* event, the lineage may traverse either the left or right parental species, according to the introgression probability (*φ* or 1 − *φ*). This probability is equivalent to the ‘inheritance probability’ *γ* of Yu *et al*. (2014) and the ‘heritability’ of Solis-Lemus and Ane (2016). The MSci model includes three sets of parameters: the speciation and introgression times (***τ***); the population size parameters (***θ***), with each *θ* = 4*Nμ*, where *N* is the effective population size and *μ* is the mutation rate per generation per site; and the introgression probabilities (***φ***). Both *τ*s and *θ*s are measured by the expected number of mutations per site. Here we assume that the MSci model is fixed; cross-model MCMC moves will be developed in future work.

Let ***G*** = {*G*_*i*_} be the set of gene trees for the *L* loci. For each locus *i, G*_*i*_ represents the gene tree topology, the branch lengths (coalescent times), and the paths taken at the *H* nodes, indicated by a set of flags for each genetree branch, with *l* for left, *r* for right and ø for null (meaning that the branch does not pass the *H* node). The data ***X*** = {*X*_*i*_} are the sequence alignments at the *L* loci. Sites within the same locus are assumed to share the same genealogical history, while the gene trees and coalescent times are assumed to be independent among loci given the species tree and parameters. The ideal data for this kind of analysis are loosely linked short genomic segments (called loci), so that recombination within a locus is unimportant while different loci are largely independent (Burgess and Yang, 2008; Hey *et al*., 2018; Lohse *et al*., 2016). The Bayesian formulation consists of two components: (i) the probability density of gene trees given the species tree under the MSci model, *f* (*G*_*i*_|***τ, θ***, ***φ***), given in Yu *et al*. (2014): Note that this density differs from that given by Kubatko (2009), as pointed out by Solis-Lemus and Ane (2016); and (ii) the likelihood of the sequence data at each locus *i* given the gene tree, *f* (*X*_*i*_|*G*_*i*_) (Felsenstein, 1981). The posterior probability density of the parameters on the species tree given sequence data is then

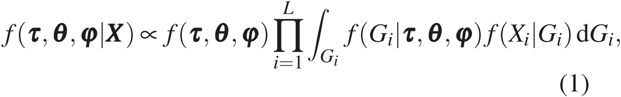

where *f* (***τ, θ***, ***φ***) is the prior on parameters. We assign inverse gamma priors on *θ*s and *τ*s and a beta prior on *φ*.

We have implemented six MCMC proposals to average over the gene trees (*G*_*i*_) and sample from the posterior (Eq. 1). Those proposals (i) change node ages on gene trees; (2) change gene tree topologies using subtree pruning and regrafting (SPR); (3) change *θ*s on the species tree using sliding windows; (4) change *τ*s on the species tree using a variant of the rubber-band algorithm (Rannala and Yang, 2003); (5) changing all node ages on the species tree and gene trees using a multiplier; and (6) change the introgression probabilities *φ*s using sliding windows. The proposals are detailed in Materials and Methods using the example species tree model of Fig. 2.

**FIG. 2.**
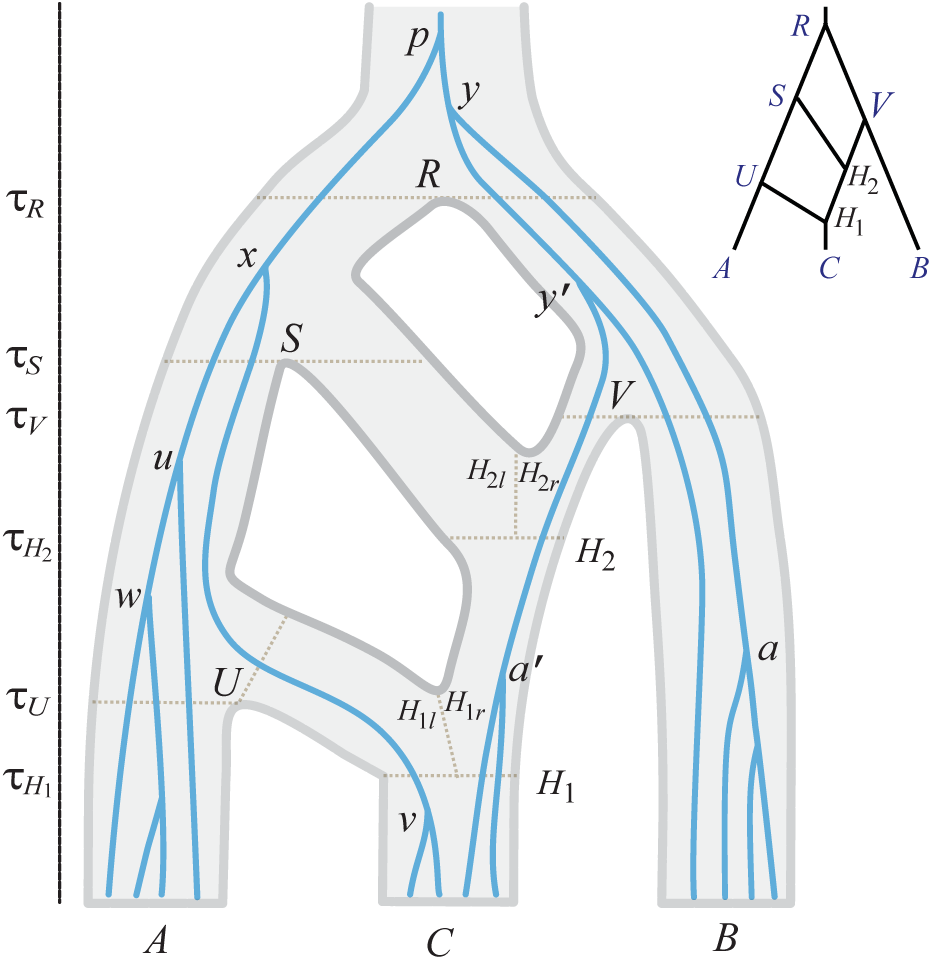
A species tree for three species (*A, B,C*) with a gene tree for 12 sequences running inside it to illustrate the gene-tree node-age move and the gene-tree SPR move. There are four speciation nodes (*R, S,U,V*) and two hybridization nodes (*H*_1_ and *H*_2_).

### Simulation study

We conducted three sets of simulation to examine the performance of BPP in different situations.

The first set includes multiple sequences from each species and examines BPP estimation of parameters in the MSci model and the impact of factors such as the number of loci, the introgression probability *φ*, and the species tree model. We used models A and C of Fig. 1. Each dataset consisted of 10, 100, or 1000 loci, with 10 sequences from each species per locus (and 30 sequences in total). We used two values for *φ* (0.1 and 0.5) and two values for *θ* (0.001 and 0.01). Either a JC (Jukes and Cantor, 1969) or GTR+Γ (Yang, 1994a, b) substitution model was used to simulate data, but JC was always used to analyze them. The results and may be summarized as follows (Figs. S1-S8).

- First, there were large variations in estimation precision and accuracy among the different parameters. For example, estimates of *θ*s for modern species were accurate even in small datasets in all combinations of trees, models, and *θ* values. In contrast *θ*s for some ancestral species (such as *θ*_*S*_, *θ*_*T*_, 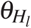 and 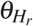 in model A when the true *θ* = 0.001) were poorly estimated, with the posterior dominated by the prior even with 100 or 1000 loci. Those parameters were hard to estimate as very few sequences enter or coalesce in the populations. These same parameters were much better estimated when the true *θ* = 0.01 as then many more sequences could enter and coalesce in the ancestral species. For similar reasons, the ages of ancestral nodes such as *τ*_*H*_ and *τ*_*S*_ were much better estimated when the true *θ* = 0.01 than when *θ* = 0.001.
- Second, parameter estimates under model C were more precise than under model A, because the former has 9 parameters while the latter 13.
- Third, there were virtually no difference in the results whether the data were simulated under JC or GTR+Γ. As the role of the mutation model in BPP is to correct for multiple hits at the same site and as the simulated sequences are highly similar, the choice of the mutation model is unimportant. Similar observations were made in previous simulations examining species tree estimation without introgression (Shi and Yang, 2018).
- Last, the data size (the number of loci) had a huge impact on the precision and accuracy of estimation. In particular, data of only 10 or 100 loci did not produce reliable estimates of *φ*, while estimates from 1000 loci were both precise (with narrow intervals) and accurate (close to the true values). Because the MSci models are parameter-rich, large datasets in the order of 1000 loci are necessary for reliable inference.

In the second set of simulations we compared BPP with two summary methods: SNAQ (Solis-Lemus and Ane, 2016; Solis-Lemus *et al*., 2017) and HYDE (Blischak *et al*., 2018), using one sequence per species. We simulated data under model A, with three ingroup species (*A, B*, and *C*), as well as two outgroup species *D* and *E*, as required by SNAQ (Solis-Lemus *et al*., 2017). One sequence was sampled per species per locus. The data were then analyzed using the three programs to estimate *φ* (Fig. S9). Data size had a large impact on the precision and accuracy of the estimates. All three methods performed poorly with 10 or 100 loci (or gene trees), but the estimates were close to the true values with 1000 loci. Overall the three methods had similar performance in estimating *φ*. In some small datasets, SNAQ and HYDE had extreme estimates of 0, while BPP always produced nonzero estimates, due to Bayesian shrinkage through the prior.

Note that the problem examined here is a conventional parameter estimation problem under a well-specified model, so that standard statistical theory applies, which states that the Bayesian method has optimal large-sample properties (O’Hagan and Forster, 2004). The small differences among the methods suggest that information in the data concerning *φ* mostly lies in the proportions of gene trees, which may be reliably estimated even if phylogenetic information content at each individual locus is low. We note that BPP has several advantages. (i) BPP accommodates the uncertainties in the data appropriately and produces posterior credible intervals (CIs), while SNAQ and HYDE generate point estimates only. (ii) BPP estimates all 13 parameters in the model while SNAQ and HYDE estimate only two (*φ* and the internal branch length) with the others unidentifiable. Estimates of ancestral population sizes (*θ*s) and species divergence and introgression times (*τ*s) may be useful for understanding the evolutionary history of the species. (iii) BPP can use loci of any data configuration, including loci with sequences from only one or two species, which are informative for BPP but carry no information about gene trees. (iv) Some introgression models or biologically important scenarios are unidentifiable using SNAQ and HYDE but can be analyzed using BPP (see below). In contrast, SNAQ and HYDE have a huge computational advantage over BPP and may be very useful for exploratory analysis in large datasets.

The third set of simulations explored the performance of BPP under models that are unidentifiable using SNAQ and HYDE. We used model D of Fig. 1 and the model (2H) of Fig. 2, with results in Fig. S10. Model D represents bidirectional introgressoin between two species. Population size parameters (*θ*s) for modern species *A* and *B* were well estimated even with 100 loci, as was *θ*_*R*_ for the root, but *θ*_*X*_ for species *X* (branch *X* – *R*) and *θ*_*Y*_ (for branch *Y*-*R*) were more poorly estimated (Fig. S10A). Both *τ* parameters were well estimated. The introgression probabilities *φ*_*X*_ and *φ*_*Y*_ were poorly estimated in small datasets of 10 or 100 loci, but were fairly accurate with 1000 loci.

Model 2H (Fig. 2) involves two introgression events on a species tree of three species. There were large differences in information content for different parameters (Fig. S10B). Parameters *θ*s for modern species were well estimated even in small datasets, but *θ*s for most ancestral species were poorly estimated because of lack of coalescent events in those populations. Parameter 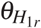 was more accurately estimated than 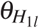 because more sequences passed node *H*_1_ from the right (with probability 1 − *φ* = 0.9) than from the left (with *φ* = 0.1), and 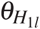 and 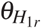 were more reliably estimated than 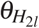 and 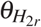 because more sequences passed node *H*_1_ than node *H*_2_. Similarly 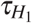 was better estimated than 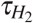. With 1000 loci, all six node ages (for *R, S,U,V, H*_1_, and *H*_2_) were well estimated. The two introgression probabilities (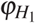 and 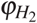) were poorly estimated with 10 or 100 loci, but were reliably estimated when 1000 loci were used.

In summary, in both introgression scenarios of models D and 2H, where SNAQ and HYDE are inapplicable, BPP appears to be a well-behaved method, providing reliable estimates of introgression probabilities as well as species divergence and introgression times.

### Analysis of the purple cone spruce data

We analyzed three datasets concerning the origin of the purple cone spruce in the Qinghai-Tibet Plateau, *Picea purpurea*, hypothesized to be a hybrid species, formed through homoploid hybridization between *P. wilsonii* (*W*) and *P. likiangensis* (*L*) (Sun *et al*., 2014). Two small datasets were previously analyzed using *BEAST under model A of Fig. 3, while the third one (the ‘Full’ data) is a much larger dataset from which the first two were sampled.

**FIG. 3.**
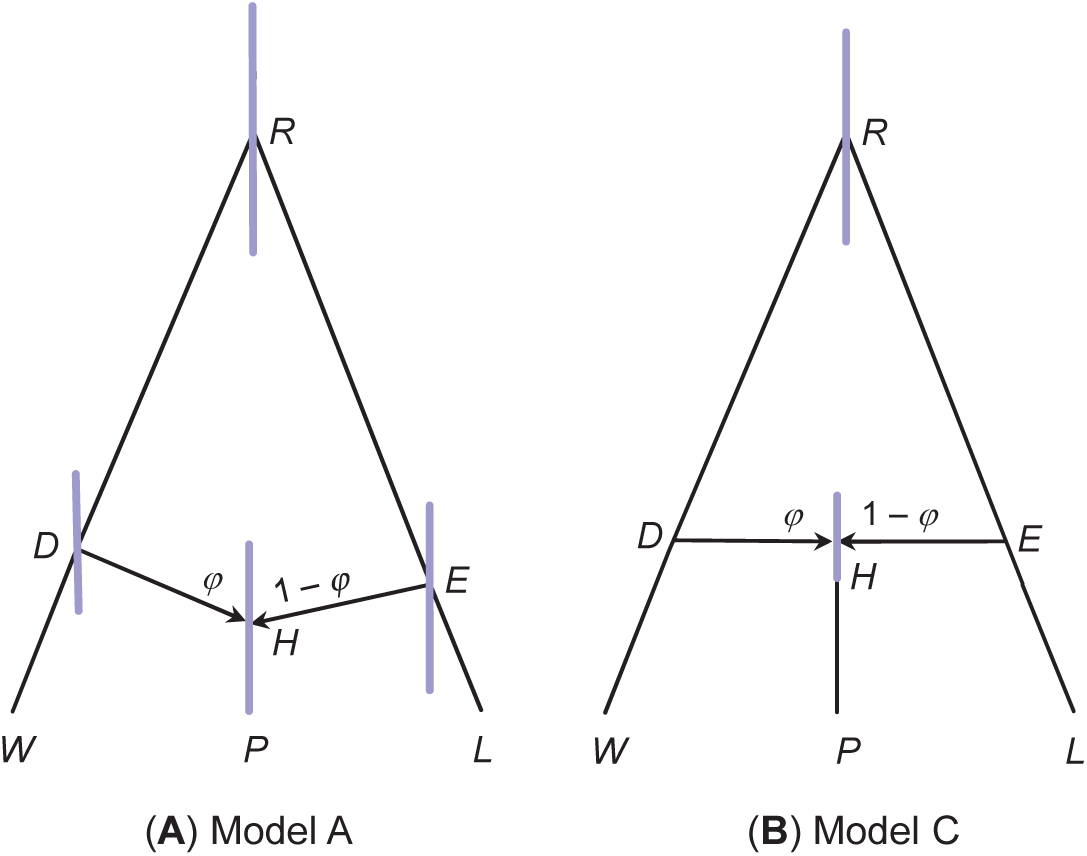
Two species trees (corresponding to models A and C of Fig. 1) for the purple cone spruce *Picea purpurea* (P) from the Qinghai-Tibet Plateau, and two parental species *P. wilsonii* (W) and *P. likiangensis* (L). The branch lengths represent the posterior means of divergence times (*τ*s) estimated from the ‘Full’ dataset, with node bars showing the 95% HPD intervals (see table S1).

Parameter estimates under model A from the two small datasets were similar to those in Zhang *et al*. (2018), with *φ* estimates between 0.32-0.44, although the estimates involved large uncertainties (table S1). The uncertainty is apparently due to the use of only 11 loci (although many sequences are available at each locus) and the shallowness of the species tree, with species divergence times being comparable with coalescent times (or with similar *τ*s and *θ*s). Accommodating rate variation among loci had very small effects. The full data produced similar parameter estimates to the two small datasets, but *φ* is larger, at 0.47-0.49. We also applied model C (Fig. 3) to the full data, which produced more precise estimates because of the smaller number of parameters (Fig. 3, table S1). The *φ* estimate under model C was 0.53 (with the 95% CI 0.36-0.71).

Note that the models represent different biological scenarios. Model A assumes the existence, and subsequent extinction at the time of hybridization, of species *DH* and *EH* (Fig. 3). This is not a very plausible model (Sun *et al*., 2014). Model C represents speciation through homoploid hybridization, with species *RDW* and *REL* coming into contact and forming a hybrid species (*H*) at time *τ*_*H*_. A possible scenario is that changes in species distribution may have led to habitat overlap between *P. wilsonii* and *P. likiangensis* during the Quaternary glaciation in the central Qinghai-Tibet Plateau (Sun *et al*., 2014). We calculated marginal likelihoods (Bayes factors) to compare models A, B, and C. The log marginal likelihood was −18,361 for model A, − 18,359 and − 18,361 for two cases of model B (with *τ*_*H*_ = *τ*_*D*_ and *τ*_*H*_ = *τ*_*E*_, respectively), and − 18,362 for model C, suggesting the fit of the models to data is similar. The marginal likelihoods are thus indecisive. We suggest that model C should be preferred, because of its biological plausibility.

### Variable introgression across the genome in the *Anopheles gambiae* species complex

To examine the variation in introgression intensity across the *Anopheles* genome, we analyzed blocks of 100 loci, assuming the species tree of Fig. 4. Estimates of *φ*_*A*→*GC*_ for the *A. arabiensis* → *A. gambiae*+*A. coluzzii* introgression and *φ*_*R*→*Q*_ for the *A. merus* → *A. quadrian-nulatus* introgression vary considerably across genomic regions or chromosomal arms (Fig. 5). The probability *φ*_*A*→*GC*_ is high (> 0.5) in most blocks, while *φ*_*R*→*Q*_ is high in 3La and 3R.

**FIG. 4.**
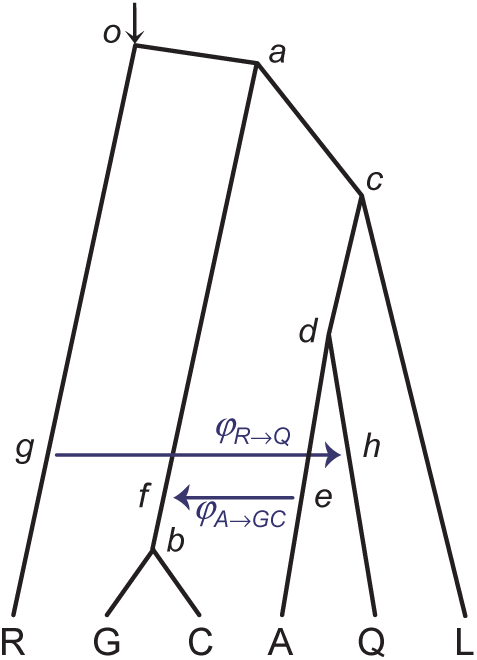
A species tree with two introgression events for the *A. gambiae* species complex.

**FIG. 5.**
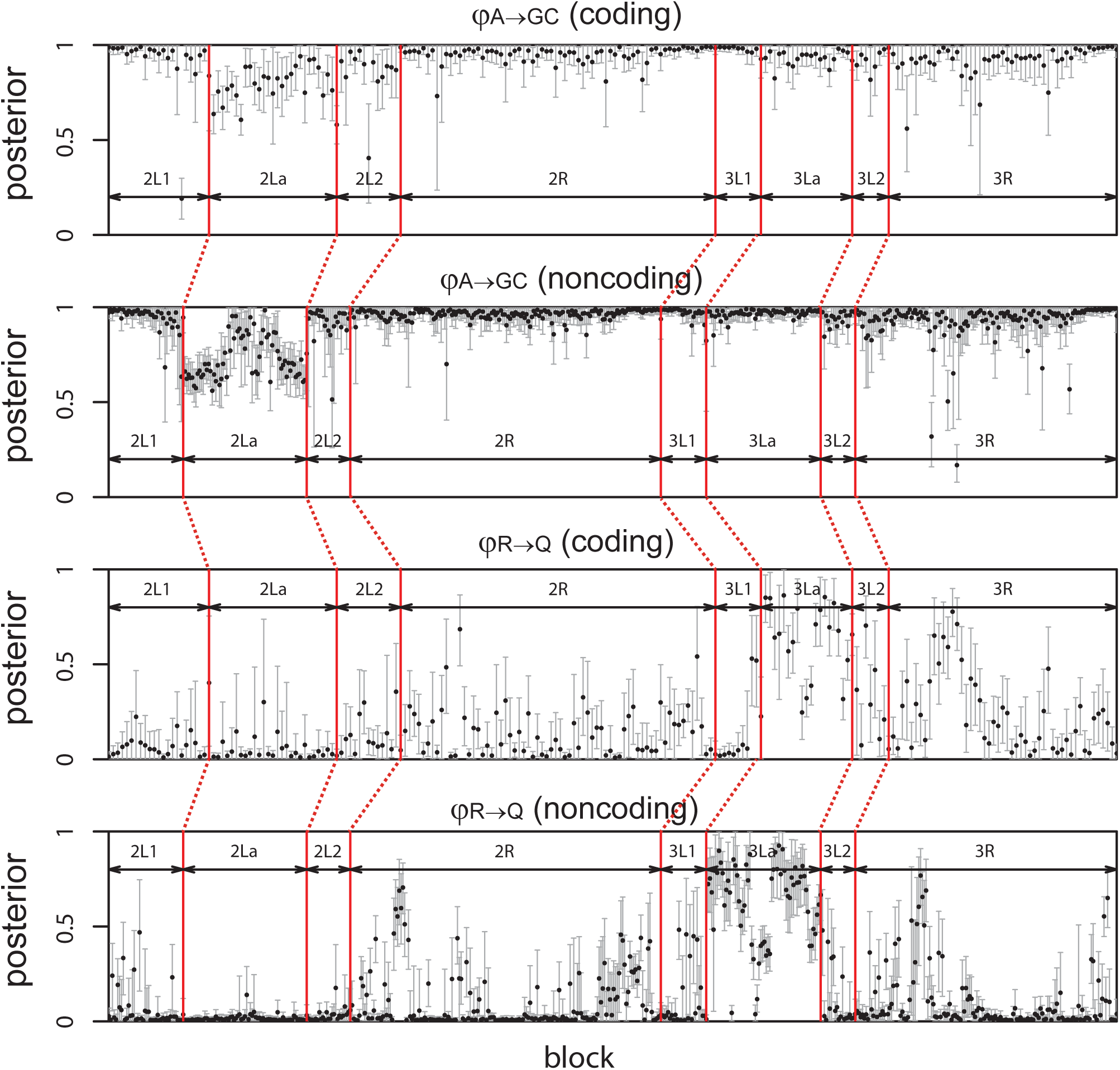
Posterior means and 95% HPD intervals of the introgression probability *φ* along the genome for the *A. gambiae* species complex, in BPP analysis of the blocks (each of 100 loci).

We then merged the loci on the same chromosomal arms/regions to form 12 large coding and noncoding datasets, and analyzed them under the model of Fig. 4 (table 1). We also sampled three sequences per locus to form data triplets for analysis using the maximum likelihood (ML) program 3S (Thawornwattana *et al*., 2018a, table S3, GAR and RQO). The introgression/migration rate from *A. arabiensis* to *A. gambiae*+*A. coluzzii* is very high (with *φ*_*A*→*GC*_ > 0.5) for all autosomes, with *M*_*A*→*G*_ ranging from 0.12 to 1.12 immigrants per generation. As noted previously (Fontaine *et al*., 2015; Thawornwattana *et al*., 2018a), the autosomes are overwhelmed by the *A* → *GC* introgression so that all species tree methods that ignore gene flow infer incorrect species trees. While in population genetic models of population subdivision, migration rates of *M* ≪ 1 do not lead to substantial population subdivision, we note that *M* as low as 0.1 may have a significant impact on the species phylogeny if the species arose through radiative speciation events and the ancestral species had large sizes.

**Table 1.** Maximum likelihood (3S) estimates of migration rate (*M* = *Nm*) and Bayesian (BPP) estimates of introgression probability (*φ*) from the *Anopheles* genomic data

| Dataset | Loci | *A. arabiensis* → *A. gambiae*+*A. coluzzii* | | | *A. merus* → *A. quadriannulatus* | | |
| --- | --- | --- | --- | --- | --- | --- | --- |
| | | $\hat{M}_{A \rightarrow G}$ (3S, GAR) | $\phi_{A \rightarrow G}$ (BPP) | | $\hat{M}_{R \rightarrow Q}$ (3S, RQO) | $\phi_{R \rightarrow Q}$ (BPP) | |
| 2L1+2 coding | 3585 | 0.319 0.372 | 0.943 (0.924, 0.963) |  | 0.025 0.020 | 0.281 (0.242, 0.322) |  |
| 2L1+2 noncoding | 6434 | 0.239 0.209 | 0.977 (0.967, 0.985) |  | 0.000 0.000 | 0.002 (0.000, 0.003) |  |
| 2La coding | 2776 | 0.915 1.120 | 0.731 (0.697, 0.766) |  | 0.052 0.036 | 0.006 (0.002, 0.009) |  |
| 2La noncoding | 6732 | 0.989 0.935 | 0.640 (0.627, 0.653) |  | 0.000 0.000 | 0.001 (0.000, 0.002) |  |
| 2R coding | 6849 | 0.511 0.477 | 0.966 (0.958, 0.975) |  | 0.030 0.024 | 0.340 (0.295, 0.391) |  |
| 2R noncoding | 17027 | 0.357 0.297 | 0.971 (0.961, 0.986) |  | 0.007 0.007 | 0.222 (0.122, 0.342) |  |
| 3L1+2 coding | 1747 | 0.168 0.340 | 0.939 (0.923, 0.956) |  | 0.000 0.024 | 0.321 (0.271, 0.369) |  |
| 3L1+2 noncoding | 4319 | 0.267 0.267 | 0.959 (0.951, 0.968) |  | 0.000 0.003 | 0.330 (0.304, 0.356) |  |
| 3La coding | 1998 | 0.866 0.790 | 0.931 (0.914, 0.947) |  | 0.127 0.105 | 0.650 (0.619, 0.680) |  |
| 3La noncoding | 6208 | 0.692 0.671 | 0.977 (0.970, 0.984) |  | 0.021 0.020 | 0.544 (0.456, 0.700) |  |
| 3R coding | 4977 | 0.393 0.365 | 0.945 (0.932, 0.958) |  | 0.042 0.042 | 0.430 (0.371, 0.488) |  |
| 3R noncoding | 14323 | 0.334 0.281 | 0.977 (0.971, 0.984) |  | 0.009 0.009 | 0.030 (0.012, 0.062) |  |

Note.— ML estimates from 3S were obtained from two random samples of the GAR and RQO triplets, while the BPP estimates (posterior means and 95% HPD intervals) used all 12 sequences at each locus.

Parameter *φ*_*R*→*Q*_ varied across chromosomal regions, and was high for the inversion region 3La. Coding and noncoding loci produced highly consistent estimates of species trees, species divergence times and population sizes (table S2) (Thawornwattana *et al*., 2018a), but estimates of migration rate/introgression probability differed between the two datasets. The high introgression rates for coding versus noncoding loci in regions 3La, 2L1+2 and 3R suggest the intriguing possibility that the introgressed genes may have brought adaptive advantages, so that introgression is aided by natural selection. The functions of coding genes or exons that are most likely transferred across the species barriers could be examined to explore this hypothesis. There is overall consistency between estimates from the IM model in 3S and the MSci model in BPP in that regions with high *φ* tend to have high *M* as well. Note that both *φ* and *M* reflect long-term effective gene flow, after the filtering of introgressed alleles by natural selection.

## Discussion

### Identifiability of MSci models

If the probability distributions of the data are identical for two sets of parameter values (Θ and Θ′), with *f* (*X*|Θ) = *f* (*X*|Θ′) for all possible data *X*, then Θ is unidentifiable given data *X*. Previous studies of identifiability have mostly focused on the use of gene tree topologies as data (Degnan, 2018; Zhu and Degnan, 2017). Note that a model unidentifiable given gene tree topologies alone may be identifiable given gene trees with branch lengths (Zhu and Degnan, 2017).

A comprehensive examination of the identifiability issue under MSci is beyond the scope of this paper. Here we consider a few simple cases. First, the population size parameter *θ* is unidentifiable if at most one sequence per locus is sampled from that species and *τ*s associated with a hybridization event may also be unidentifiable. Consider the model of Fig. 2 and suppose the data consist of one sequence from each species. Then *θ*_*A*_, *θ*_*B*_ and *θ*_*C*_, as well as 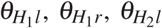 and 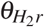, are unidentifiable. In addition, 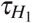 and 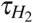 are unidentifiable. Parameters *τ*_*U*_, *τ*_*V*_, *τ*_*S*_, *τ*_*R*_, and 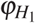 and 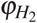 are identifiable, as are *θ*_*R*_, *θ*_*S*_, *θ*_*U*_, and *θ*_*V*_. In this case, the gene tree at any locus depends on whether sequence *c* takes the left path at *H*_1_ and enters species *U* (which happens with probability 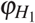), or it takes the right path and enters species *H*_2_ (which happens with probability 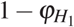), but not on the age of the *H*_1_ node. The same applies to the path taken by sequence *c* at *H*_2_.

The most interesting case for the MSci model implemented here is where multiple sequences are sampled from each species at each locus, with multiple sites per locus. We speculate that the MSci model is identifiable on such data of sequence alignments as long as it is identifiable when the data consist of gene trees with coalescent times: Θ is identifiable using multi-locus data *X* if and only if *f* (*G*, ***t***|Θ) ≠ *f* (*G*, ***t***|Θ′) for some *G* and ***t***. Note that identifiability implies statistical consistency for a full-likelihood method as implemented here. If the model is identifiable, the Bayesian parameter estimates will approach the true values when the number of loci approaches infinity.

Here we note an interesting unidentifiability issue with model D of Fig. 1. Let Θ = (*θ*_*A*_, *θ*_*B*_, *θ*_*R*_, *θ*_*X*_, *θ*_*Y*_, *τ*_*R*_, *τ*_*X*_, *φ*_*X*_, *φ*_*Y*_) be the parameters of the model, and let Θ′ have the same parameter values as Θ except that 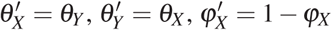, and 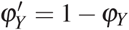. Then *f* (*G*|Θ) = *f* (*G*|Θ′) for any Θ, *G*, and data configuration (with *n*_*A*_ and *n*_*B*_ sequences from *A* and *B*, respectively, say). Thus for every point Θ in the parameter space, there is a ‘mirror’ point Θ′ with exactly the same likelihood. With Θ, a certain number of *A* sequences may take the left (upper) path at *X* (with probability *φ*_*X*_) and enter population *XR*, coalescing at the rate 2*/θ*_*X*_, while with Θ′, the same *A* sequences may take the right (horizontal) path (with probability 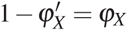) and enter population *Y R*, coalescing at the rate 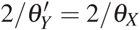. The differences between the two scenarios are in the labelling only, with ‘left’ and *X* under Θ corresponding to ‘right’ and *Y* under Θ′, but the probabilities involved are exactly the same. The same argument applies to sequences from *B* going through node *Y*, and to sequences from *A* and *B* considered jointly. This is a case of the label-switching problem. Arguably Θ and Θ′ have the same biological interpretations concerning the relatedness of the sequences sampled from *A* and *B*. If the priors on *φ*_*X*_ and *φ*_*Y*_ are symmetrical, say beta(*α, α*), the posterior density will satisfy *f* (Θ|***X***) = *f* (Θ′|***X***) for all Θ. Otherwise the “twin towers” may not have exactly the same height.

Note that the label-switching kind of unidentifiability does not hinder the utility of the model. One can apply an identifiability constraint, such as *φ* < 0.5, to remove the unidentifiability. However, in the general case of multiple bidirectional introgression events or multiple species on the species tree, it may be complicated to decide on the identifiability of the model.

Finally we point out that there are many scenarios of data configurations and parameter settings in which some parameters are only weakly identifiable and very hard to estimate. For example, if *θ*_*C*_ is very small relative to 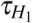 in Fig. 2, sequences from *C* will have coalesced before reaching node *H*_1_, so that only one *C* sequence passes *H*_1_ and the data will have little information about 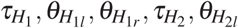, and 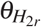.

### Full-likelihood and summary methods to accommodate introgression/migration

While biologically simplistic, the MSci and IM models offer powerful tools for analysis of genomic sequence data from closely related species, when cross-species gene flow appears to be the norm (Mallet *et al*., 2016; Martin and Jiggins, 2017). Full likelihood implementations of those models, including the ML (Dalquen *et al*., 2017; Zhu and Yang, 2012) and Bayesian MCMC methods (Hey *et al*., 2018; Wen and Nakhleh, 2018; Zhang *et al*., 2018), make efficient use of the information in the data and naturally accommodate phylogenetic uncertainties at individual loci caused by high sequence similarities (Edwards *et al*., 2016; Xu and Yang, 2016). The complexity of those models means that large datasets with hundreds or thousands of loci may be necessary to obtain reliable parameter estimates, as indicated by our analyses of both simulated and real data. We have implemented several MSci models, of types A, B, C and D of Fig. 1, and applied them to large datasets of over 10,000 loci (table 1). We suggest that this is a promising start, from which further improvements to the mixing efficiency may be possible. Future work will include implementation of MCMC proposals to move between MSci models, and systematic examination of the identifiability issues.

Our simulation suggests that under simple introgression scenarios, summary methods such as SNAQ and HYDE can produce as reliable estimates of the introgression probability as BPP. However, full likelihood methods provide measures of uncertainties, and are applicable to complex introgression scenarios which are unidentifiable using summary methods. Summary methods are simple to implement, computationally efficient, and useful for analyzing large datasets. They can be used to generate hypotheses for further testing and estimation using BPP. Furthermore, the current implementation of MSci in BPP assumes the molecular clock and is unsuitable for distantly related species. Summary methods such as SNAQ use outgroups to root the tree without the need for the molecular clock.

### Variable introgression probability across the genome

The models implemented here (Fig. 1A-C) assume that the introgression probability or migration rate is constant among loci or across the genome. However, the impact of introgressed alleles on the fitness of the individual may strongly depend on the function of the genes in the introgressed region. Genes involved in cross-species incompatibilities are unlikely to be accepted in the recipient species. For example, crossing experiments between *A. arabiensis* and *A. gambiae* highlighted large differences between the chromosomes, with the X chromosome being most resistant to introgression, presumably because it harbors genes involved in cross-species sterility and inviability (Slotman *et al*., 2005). Differential selection across the genome means that the *φ* parameter should vary among loci. Note that *φ* in our models when estimated from genetic sequence data reflects the long-term combined effects of migration, recombination and natural selection. It may be very different from the per-generation hybridization rate, which should apply to the whole genome.

In our analysis of the *Anopheles* genomic data, we used blocks of 100 loci to partially accommodate the variation in migration rate or introgression probability across chromosomal regions (Fig. 5). We leave it to future work to implement MSci models with *φ* varying among loci. We note that many sequences per locus may be necessary to estimate locus-specific migration rates or introgression probabilities.

## Materials and Methods

### MCMC proposals

We have adapted the five proposals in Rannala and Yang (2003) to accommodate hybridization nodes on the species tree, and added another move to update the *φ* parameters.

Step 1. Change node ages on gene trees using a sliding window. Suppose the concerned gene-tree node is node *x* with age *t* in population *X*, with parent node *p* in population *P* and two daughter nodes *u* and *v* in populations *U* and *V*, respectively. To propose a new node age 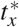 first determine the bounds, 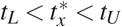, with *tU* determined by the age of the parent node (*t*_*p*_) and *t*_*L*_ by the age of the oldest daughter node: *t*_*L*_ ≥ max(*t*_*u*_, *t*_*v*_). In addition, if the two daughter nodes are in different populations (with *U* ≠ *V*), *t*_*L*_ must be older than the youngest common ancestor of populations *U* and *V* on the species tree.

Generate the new age 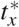 by sampling around *t*_*x*_, reflected into the interval (*t*_*L*_, *t*_*U*_). The new node *x*\* has to reside in a population that is descendant to the mother population *P* and ancestral to the daughter populations *U* and *V*. Among those target populations, we sample one uniformly. Given the sampled population for *x*\*, we sample the flags for the three branches: *p*–*x*\*, *x*\*– *u*, and *x*\*–*v*. In each case the two ends of the branch are already assigned a population. This move may cause large changes to the flags even though it does not change the gene-tree topology. For example, consider the change of *t*_*x*_ in Fig. 2. Node *x* is in population *S*, with branch *x*– *v* having the flags *l*ø, since the branch passes *H*_1_ from the left and does not pass *H*_2_. Suppose the new age, generated in the interval (*t*_*u*_, *t*_*p*_), is 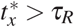 so that the new node *x*\* resides in *R*. The re-sampled flags for branch *x*\*–*v* may be *rr*, if the new branch passes both *H*_1_ and *H*_2_ from the right. The proposal ratio is given by the probabilities of sampling the flags at the *H* nodes.

Step 2. Subtree pruning and regrafting (SPR) move to change the gene tree topology. This move cycles through the non-root nodes on the gene trees. Suppose the node is *a*. We prune off its mother node *y*. The remaining part of the gene tree is called the backbone. We sample the new age 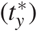 before reattaching the subtree *y*–*a* onto the backbone. This move always changes the node age *t*_*y*_ but may not change the gene tree topology.

First we determine the bounds on the age of reattachment point: 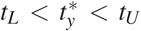. The maximum age is unbounded, while the minimum is *t*_*L*_ ≥ *t*_*a*_. However, if there are no branches on the backbone passing the population of node *a*, the reattachment point has to be in an ancestral species (in which there exists at least one branch on the backbone) and *t*_*L*_ has to be greater. For example, clade *a* in Fig. 2 resides in population *B*, and if we prune off clade *a*, there will still be branches in *B* on the backbone for reattachment. In contrast, clade *a*′ resides in population *H*_1*r*_, but if we prune off *y*′-*a*′, there will be no branches in population *H*_1*r*_ for reattachment and the youngest ancestor of *H*_1*r*_ with branches on the backbone is *V*, so that *t*_*L*_ = *τ*_*V*_.

We generate a new age 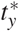 around the current age (*t*_*y*_), reflected into the interval (*t*_*L*_, *t*_*U*_) if necessary, and then reattach *y* and clade *a* to a branch on the backbone at time 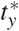. A feasible target branch should cover 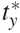, and should at time 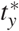 be in a population ancestral to the population of *a*. We sample a target branch at random, either uniformly or with weights determined using local likelihoods.

In case the new branch *y*\*-*a* passes hybridization nodes, we sample the flags at each hybridization node, as in step 1. Suppose we prune off branch *y*′–*a*′ in Fig. 2 and the new age is 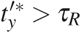. Then we let branch *y*′*–*a*′ go through *H*_2*l*_ or *H*_2*r*_ according to their probabilities. The proposal ratio is given by the number of target branches for reattachment and the probabilities for sampling the flags.

Step 3. Change *θ*s on the species tree using a sliding window. This step is the same as in Rannala and Yang (2003).

Step 4. Change *τ*s on the species tree using a variant of the rubber-band algorithm (Rannala and Yang, 2003). We generate a new age (*τ*\*) around the current age, reflected into the interval (*τ*_*L*_, *τ*_*U*_), determined using the ages of the parent nodes and daughter nodes on the species tree. Next we change the ages of the affected nodes on the gene trees using the rubber-band algorithm. An affected node has age in the interval (*τ*_*L*_, *τ*_*U*_) and resides in the current population (with age *τ*) or the two daughter populations (if a speciation node is changed), or in the two current populations (*H*_*l*_ and *H*_*r*_) and the daughter population (if an *H* node is changed). For example, to change *τ*_*S*_, the bounds are 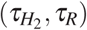, and the affected nodes on the gene tree of Fig. 2 are in species *S, U* and *H*_2_. These are *x, u*, and *w*. To change 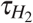 the bounds are 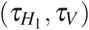 and the affected nodes are in species *H*_2*l*_, *H*_2*r*_, and *H*_1*r*_, and are *a*′. The proposal for changing node ages on the gene tree given the bounds is as in Rannala and Yang (2003, Eqns A7 and A8).

Step 5. Rescale all node ages on the species tree and on the gene trees using a mixing step (a multiplier) (Rannala and Yang, 2003).

Step 6. Change the introgression probability *φ* for each introgression event using a sliding window. This step affects the gene-tree density, but not the sequence likelihood.

The sliding window used in BPP is the Bactrian move with the triangle kernel (Thawornwattana *et al*., 2018b; Yang and Rodriguez, 2013). Step lengths are adjusted automatically during the burnin, to achieve an acceptance rate of ∼ 30% (Yang and Rodriguez, 2013).

### Simulation study

We conducted three sets of simulations. The first set includes multiple sequences from each species and examines BPP estimation of parameters in the MSci model and the impact of the number of loci, the introgression probability *φ*, and the species tree model. The second set compares BPP with two summary methods: SNAQ (Solis-Lemus and Ane, 2016; Solis-Lemus *et al*., 2017) and HYDE (Blischak *et al*., 2018), using one sequence per species. The third set explores the performance of BPP when the model is unidentifiable using SNAQ and HYDE.

For the first set of simulations, multi-locus datasets were simulated under the MSci models A and C of Fig. 1 and then analyzed using BPP to examine the precision and accuracy of parameter estimation. For model A we used *τ*_*R*_ = 0.03, *τ*_*S*_ = 0.02, *τ*_*T*_ = 0.02, and *τ*_*H*_ = 0.01. For model C we used *τ*_*R*_ = 0.03 and *τ*_*S*_ = *τ*_*T*_ = *τ*_*H*_ = 0.01. We used two values of *φ* (0.1 and 0.5) and two values of *θ* (0.001 and 0.01), applied to all populations. Each dataset consisted of 10, 100 or 1000 loci, and at each locus 10 sequences were sampled from each species (with 30 sequences in total). The sequence length was 500 sites.

Data were generated using the ‘simulate’ option of BPP. Gene trees with branch lengths (coalescent times) were simulated under the MSci model. Then sequences were “evolved” along the branches of the gene tree according to either the JC (Jukes and Cantor, 1969) or GTR+Γ (Yang, 1994a, b) models, and the sequences at the tips of the gene tree constituted the data at the locus. In the GTR+Γ model, the GTR parameters varied among loci according to estimates obtained for chromosomal arm 2L from the *Anopheles gambiae* species complex (Thawornwattana *et al*., 2018a). The base-frequency parameters were generated from a Dirichlet distribution (*π*_*T*_, *π*_*C*_, *π*_*A*_, *π*_*G*_) ∼ Dir(25.18, 20.50, 25.22, 20.38). The GTR exchangeability parameters (Yang, 1994a) were (*a, b, c, d, e, f*) ∼ Dir(7.59, 3.23, 2.95, 2.93, 2.93, 7.57). The overall rates for loci varied according to a gamma distribution *G*(5, 5), while the rates for sites at the same locus varied according to the gamma distribution with mean one, *G*(*α, α*) (Yang, 1994b), with the shape parameter *α* sampled from *G*(20, 4).

The number of replicates was 10. Thus with 2 trees (*A* and *C*), 2 *φ* values (0.1 and 0.5), 2 *θ* values (0.001 and 0.01), 2 mutation models (JC and GTR+Γ), and 3 data sizes (10, 100, and 1000 loci), a total of 480 = 2 × 2 × 2 × 2 × 3 × 10 replicate datasets were generated.

Each dataset was analyzed using BPP. The JC model was always assumed whether the data were simulated under JC or GTR+Γ. Inverse-gamma priors were assigned on parameters *θ* and *τ*_0_ (the root age), with the shape parameter 3 and the prior mean equal to the true value: IG(3, 0.02) for *θ* = 0.01 and IG(3, 0.002) for *θ* = 0.001, and *τ*_0_ ∼ IG(3, 0.06). The inversegamma distribution with shape parameter *α* = 3 has the coefficient of variation 1 and constitutes a diffuse prior. The uniform prior 𝕌(0, 1) was used for *φ*.

Pilot runs were used to determine the suitable chain length, and then the same settings (such as the burn-in, the number of MCMC iterations, and the sampling frequency) were used to analyze all replicates. Convergence was assessed by running the same analysis multiple times and confirming consistency between runs (Flouris *et al*., 2018; Yang, 2015).

The second set of simulation was to compare BPP with summary methods. Most methods are designed to test for the presence of gene flow (hybridization or migration) (Degnan, 2018). Here we used two methods that can estimate the introgression probability under a fixed introgression model: SNAQ (Solis-Lemus and Ane, 2016) implemented in the program PhyloNetworks (Solis-Lemus *et al*., 2017) and HYDE (Blischak *et al*., 2018). The basic algorithms for SNAQ and HYDE are formulated for the case of three species with one or two outgroup species used to root the tree. SNAQ uses the proportions of the three gene tree topologies, based on the observation that the probabilities for the two mismatching gene trees (which have different topologies from the species tree) are the same if there is deep coalescent but no gene flow while they are different if there is gene flow as well (Yu *et al*., 2014). HYDE uses the proportions of the three parsimony-informative site patterns pooled across loci or genomic regions (*xxyy, xyxy* and *xyyx*), based on the observation that the probabilities for the two “mismatching” site patterns (*xyxy* and *xyyx*) are the same if there is deep coalescent but no gene flow while these are different if there is gene flow as well (Green *et al*., 2010).

We used model A of Fig. 1, plus two outgroup species *D* and *E*, to simulate *L* = 10, 100 or 1000 loci, with one sequence per species per locus. The data were then analyzed using SNAQ and HYDE, as well as BPP. The JC model was used both to simulate and to analyze the data. For SNAQ, gene trees were inferred using RAxML (Stamatakis *et al*., 2012). For BPP, the point estimates (posterior means) of *φ* were used for comparison even though it produced estimates for all parameters, with CIs.

The third set of simulation explores the performance of BPP under models that are unidentifiable using SNAQ and HYDE. An MSci model may be identifiable given the gene trees with coalescent times but unidentifiable given gene tree topologies only (Degnan, 2018). We simulated and analyed data using BPP under two models: model D of Fig. 1 with bidirectional introgression between species *A* and *B* and the model of Fig. 2 (referred to as model 2H), with three species and two introgression events. Under model D, there is only one gene tree between two species so that its frequency is uninformative and SNAQ is not applicable, and nor is HYDE. Under model 2H, frequencies of three gene trees or three site patterns cannot be used to estimate two introgression probabilities and two internal branch lengths: it is thus impossible to apply SNAQ and HYDE to such data.

For model D, we used the following parameter values: *τ*_*R*_ = 0.01, *τ*_*X*_ = *τ*_*Y*_ = 0.005, *φ*_*X*_ = 0.1, *φ*_*Y*_ = 0.3, and *θ* = 0.01 for all populations. We simulated 10 replicate datasets, each of 10, 100, or 1000 loci. At each locus, we sampled 10 sequences per species (20 sequences in total), with the sequence length to be 500. The JC mutation model was used both to simulate and to analyze data by BPP. Note that there is an interesting identifiability issue (or label-switching issue) with model D, such that the two sets of parameters Θ = (*θ*_*A*_, *θ*_*B*_, *θ*_*R*_, *θ*_*X*_, *θ*_*Y*_, *τ*_*R*_, *τ*_*X*_, *φ*_*X*_, *φ*_*Y*_) and Θ′ = (*θ*_*A*_, *θ*_*B*_, *θ*_*R*_, *θ*_*Y*_, *θ*_*X*_, *τ*_*R*_, *τ*_*X*_, 1 − *φ*_*X*_, 1 − *φ*_*Y*_) are unidentifiable (see Discussion). Thus an identifiability constraint should be applied, such as *φ*_*X*_ < 0.5. We ran the MCMC without any constraint, but the MCMC sample was pre-processed, with Θ replaced by Θ′ if the sampled value for *φ*_*X*_ > 0.5, before the posterior summary was generated.

For model 2H (Fig. 2), the following parameter values were used: 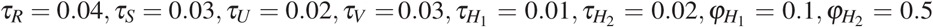 and *θ* = 0.01 for all populations. As above, 10 sequences per species were generated, with 30 sequences per locus. The sequence length was 500. The JC model was used both to simulate and to analyze the data.

### Analysis of the purple cone spruce datasets

We reanalyzed sequence data concerning the origin of the purple cone spruce *Picea purpurea* (P) from the Qinghai-Tibet Plateau, hypothesized to have originated through homoploid hybridization between *P. wilsonii* (W) and *P. likiangensis* (L) (Sun *et al*., 2014). The data were generated by Sun *et al*. (2014). To make the computation feasible for the *BEAST program, Zhang *et al*. (2018) constructed and analyzed two non-overlapping data subsets (datasets 1 and 2), each with 40, 30, and 30 phased sequences for P, W, and L, respectively, at 11 autosomal loci. We analyzed these datasets for comparison with the analysis of Zhang *et al*. (2018) using *BEAST. We were unsuccessful to apply PHYLONET (Wen and Nakhleh, 2018) to analyze those datasets, which appeared to be too large for the program. We also used BPP to analyze the ‘Full’ dataset from which datasets 1 and 2 were sampled, with 112, 100, and 120 sequences per locus for the same 11 loci.

The species tree of Fig. 3 was assumed (Sun *et al*., 2014). The priors were *τ*_0_ ∼ IG(3, 0.004), *θ* ∼ IG(3, 0.003), and *φ* ∼ 𝕌(0, 1). Rates for loci were either constant or had a Dirichlet distribution with *α* = 2 (Burgess and Yang, 2008). We used a burn-in of 32,000 iterations and took 10^5^ samples, sampling every 10 iterations. The program was run at least twice for each analysis, to check for consistency between runs. Each run (on a single core) took ∼ 5 days.

Marginal likelihood for models A, B, and C (Fig. 1) were calculated using thermodynamic integration with Gaussian quadrature (Lartillot and Philippe, 2006; Rannala and Yang, 2017), with 16 quadrature points.

### Analysis of the genomic data from the *Anopheles gambiae* species complex

We used the coding and noncoding loci compiled by Thawornwattana *et al*. (2018a) from the genomic sequences for six species in the *A. gambiae* species complex: *A. gambiae* (G), *A. coluzzii* (C), *A. arabiensis* (A), *A. melas* (L), *A. merus* (R), and *A. quadriannulatus* (Q) (Fontaine *et al*., 2015). There are 12 sequences per locus, with two sequences per species.

We analyzed blocks of 100 loci, as in Thawornwattana *et al*. (2018a), and then combined loci for each of the eight chromosomal arms/regions: 2L1, 2La (the inversion region on 2L), 2L2, 2R, 3L1, 3La (the inversion region on 3L), 3L2, and 3R. Since our objective was to estimate the introgression probability for the autosomes, the X chromosome was not used. The species tree is in Fig. 4, from Thawornwattana *et al*. (2018a, fig. 6).

The priors were *τ*_0_ ∼ IG(3, 0.2) with mean 0.1 for the age of the root, *θ* ∼ IG(3, 0.04) with mean 0.02, and *φ* ∼ 𝕌(0, 1). We used a burn-in of 16,000 iterations, and took 5 × 10^5^ samples, sampling every 2 iterations. Pilot runs suggest that this generates effective sample size (ESS) > 1000. Each analysis of the block took a few hours while the analysis of the 12 large combined datasets of table 1 each took 1-2 weeks.

For comparison, we used the ML program 3S (Dalquen *et al*., 2017; Zhu and Yang, 2012) to estimate the migration rate *M* = *Nm* under the IM model. The implementation assumes three species (1, 2, and 3, say), with three sequences per locus. We sampled three sequences, with half of the loci having the ‘123’ configuration, a quarter with ‘113’, and another quarter with ‘223’ (Thawornwattana *et al*., 2018a). We generated two replicate datasets by sampling the GAR and RQO triplets to estimate the migration rate *M*_*A*→*G*_ and *M*_*R*→*Q*_ (Fig. 4). While limited to three sequences, 3S can use tens of thousands of loci and each run took a few minutes.

## Software Availability

The algorithms described in this article are implemented in the program BPP Version 4 (Flouris *et al*., 2018; Yang, 2015), which may be downloaded from https://github.com/BPP. The python3 code and scripts for simulating and analyzing the sequence data and for making plots using ggplot are at the github site https://github.com/brannala/NetworkMSCSimulations.

## Acknowledgments

This study has been supported by Biotechnological and Biological Sciences Research Council grants (BB/N000609/1 and BB/P006493/1) to Z.Y.

**Appendix A. Extended Newick notation for the MSci model**

We use the extended Newick notation (Cardona *et al*., 2008) to represent the MSci model in the BPP program. The parenthesis notation ‘(*A, B*)*S*’ specifies two branches from the speciation node *S* to two daughter species *A* and *B*, while ‘(*A*)*H*’ specifies one branch from *H* to *A*. Every branch is represented once. Each tip species occurs once. Internal nodes for speciation nodes may and may not be labeled, but hybridization (*H*) nodes must be labeled. In models A-C of Fig. 1, each *H* node occurs twice in the notation, once as a label for an ancestral node and another time as a tip, and the introgression probability *φ* is identified with the ancestral node (while its ‘mirror’ tip node has 1 − *φ*). Thus models A-C of Fig. 1 are represented as ((*A*, (*C*)*H*)*S*, (*H, B*)*T*) *R*, with parameter *φ* assigned to the *SH* branch and 1 − *φ* to the *TH* branch. The extended Newick notation is not unique. The representation ((*A, H*)*S*, ((*C*)*H, B*)*T*) *R* specifies an equivalent model, with parameter *φ* assigned to branch *TH* and 1 − *φ* to branch *SH*.

The three types of models in Fig. 1 (A, B and C) are distinguished using the metadata variable ‘tau-parent’, which is assigned the value ‘yes’ or ‘no’ depending on whether the parent node has an age (*τ*) distinct from that of the hybridization node. Thus models A, B, and C of Fig. 1 are represented as

(A): ((*A*, (*C*)*H*[&tau-parent = yes])*S*, (*H*[&tau-parent = yes], *B*)*T*) *R*.

(B): ((*A*, (*C*)*H*[&tau-parent = no])*S*, (*H*[&tau-parent = yes], *B*)*T*) *R*.

(c): ((*A*, (*C*)*H*[&tau-parent = no])*S*, (*H*[&tau-parent = no], *B*)*T*) *R*.

Model D (bidirectional introgression) differs from models A-C in that each of nodes *X* and *Y* has two parent nodes and two daughter nodes. The model is represented as ((*A*, (*B*)*Y*) *X*, (*X*) *Y*) *R*. The notation is again not unique, and an equivalent representation is (((*A*)*X*, *B*)*Y*, (*Y*) *X*) *R*. In both notations, the *φ* parameter is assigned to the branch with an older parent while the horizontal branch has 1 − *φ*.

As a more complex example, the species graph for the *Anopheles* mosquitoes of Fig. 4 is represented as ((*R*, (*Q*)*h*[&tau-parent = no])*g*, (*f* [&tau-parent = yes], (((((*G,C*)*b*) *f* [&tau-parent = no], *A*)*e, h*[&tau-parent = yes]) *d, L*)*c*)*a*)*o*.

## Supplementary material

**Figure S1:**
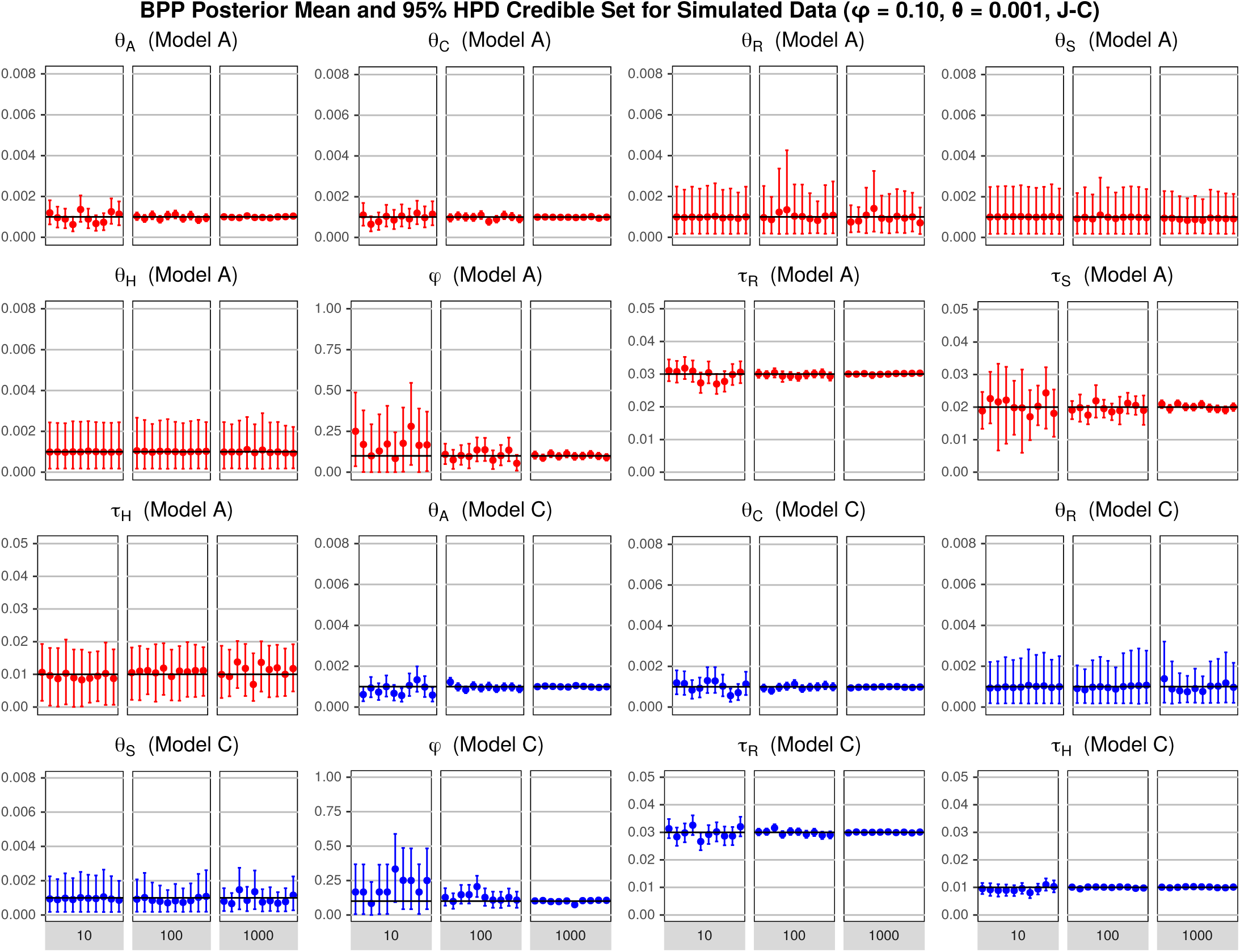
Posterior means and 95% HPD credibility intervals (CI) of parameters under models A and C of Fig. 1 in 10 replicate datasets, each of 10, 100, or 1000 loci, simulated under the parameter combinations: *φ* = 0.1, *θ* = 0.001, and the JC mutation model. The horizontal line represents the true parameter value. Note that there are 13 parameters under model A 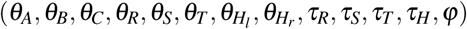, and 9 under model C (*θ*_*A*_, *θ*_*B*_, *θ*_*C*_, *θ*_*R*_, *θ*_*S*_, *θ*_*T*_, *τ*_*R*_, *τ*_*S*_, *φ*). Due to the symmetry of the experimental design, the results for some parameters are identical so that only the non-redundant results are shown to save space: for example, *θ*_*A*_ is shown but *θ*_*B*_ is not.

**Figure S2:**
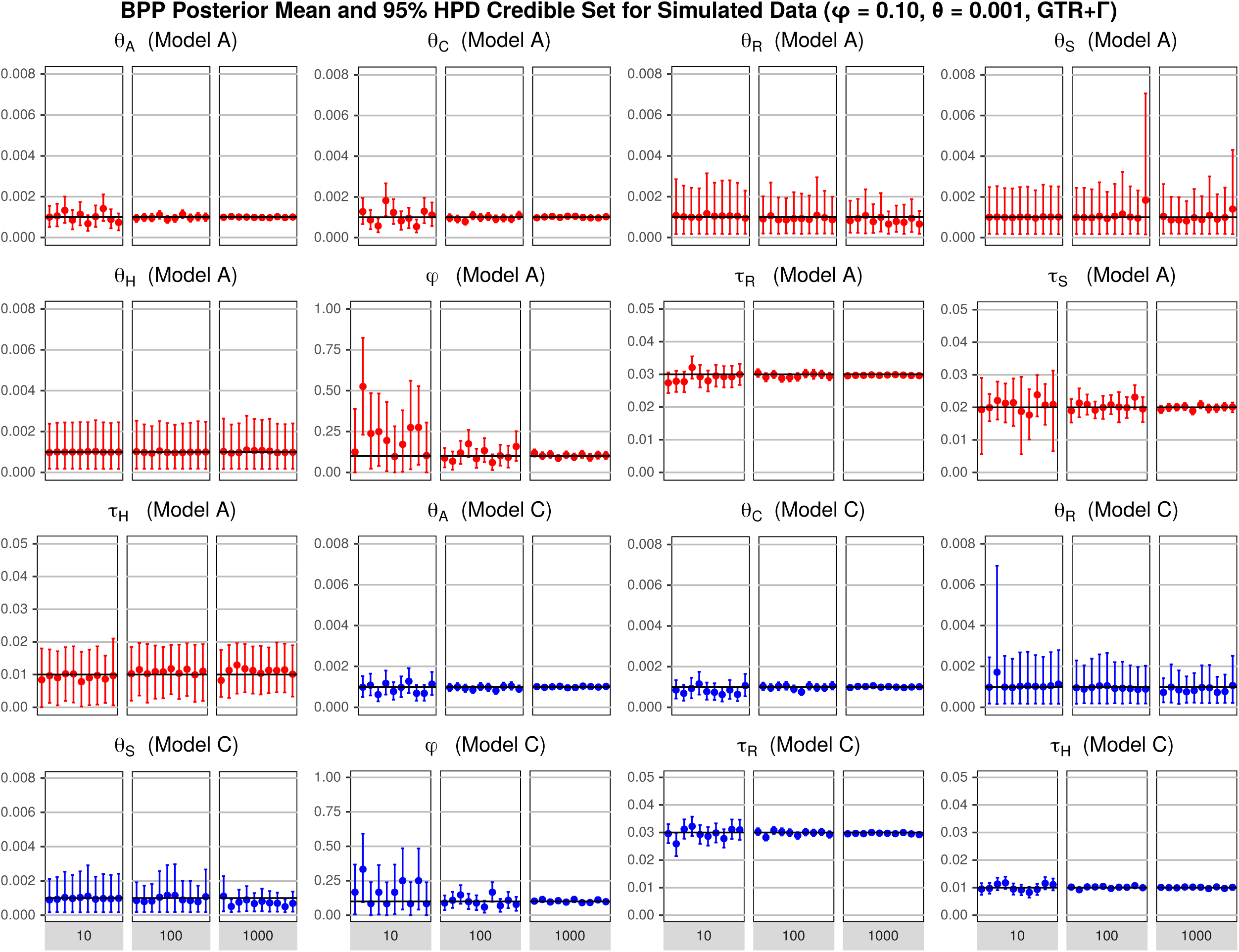
Simulation results for the combination *φ* = 0.1, *θ* = 0.001, and the GTR+Γ mutation model. See legend to Fig. S1.

**Figure S3:**
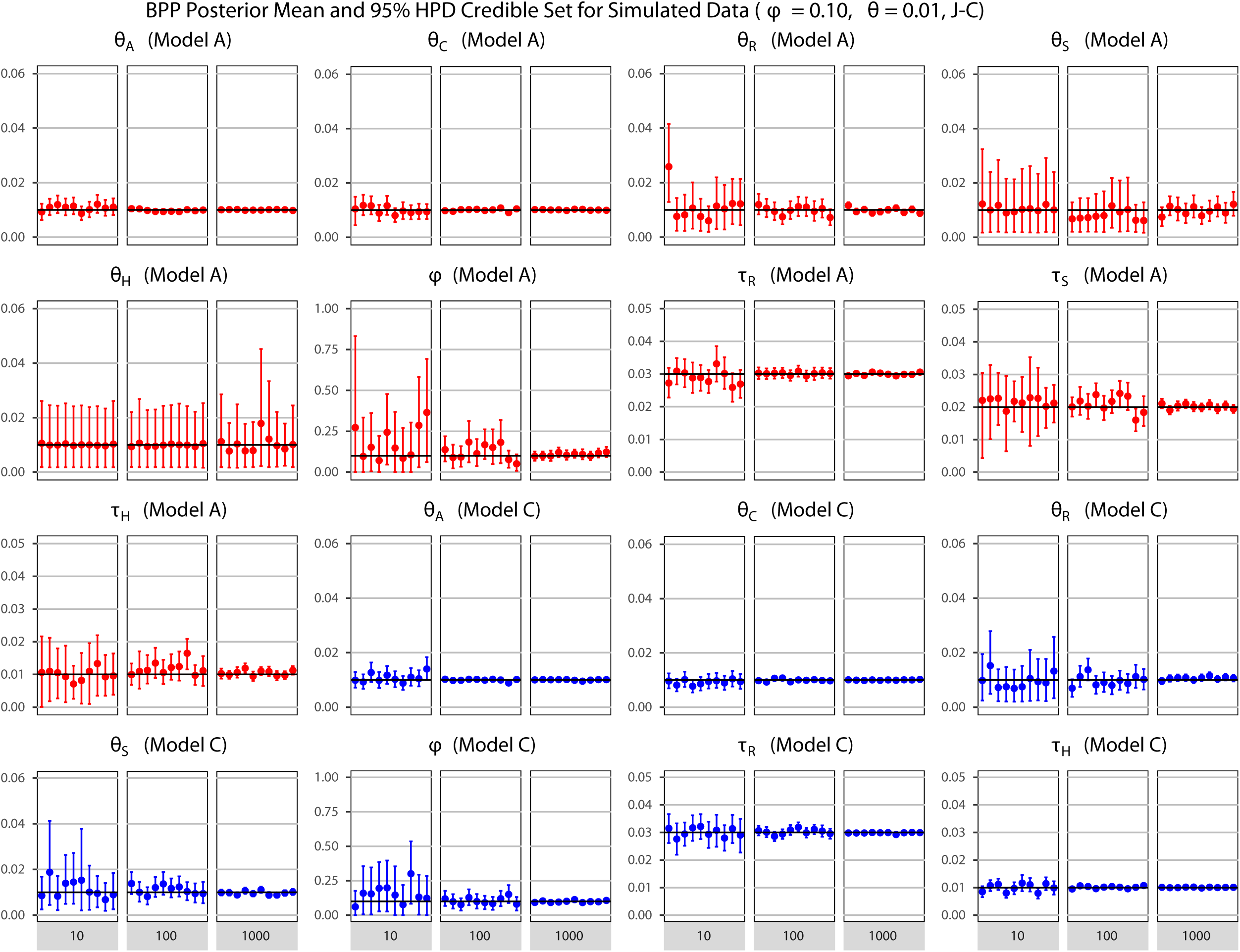
Simulation results for the combination *φ* = 0.1, *θ* = 0.01, and the JC mutation model. See legend to Fig. S1.

**Figure S4:**
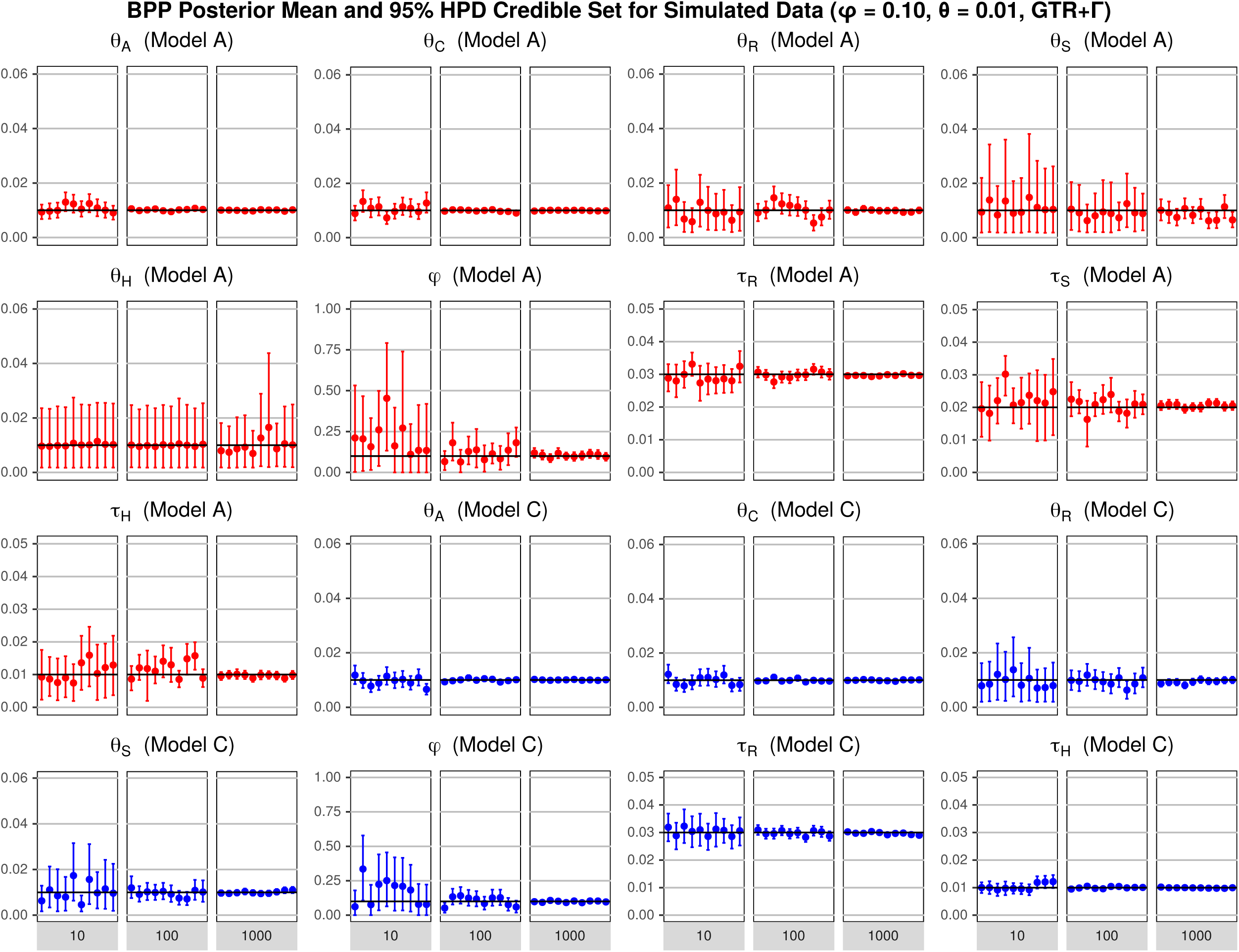
Simulation results for the combination *φ* = 0.1, *θ* = 0.01, and the GTR+Γ mutation model. See legend to Fig. S1.

**Figure S5:**
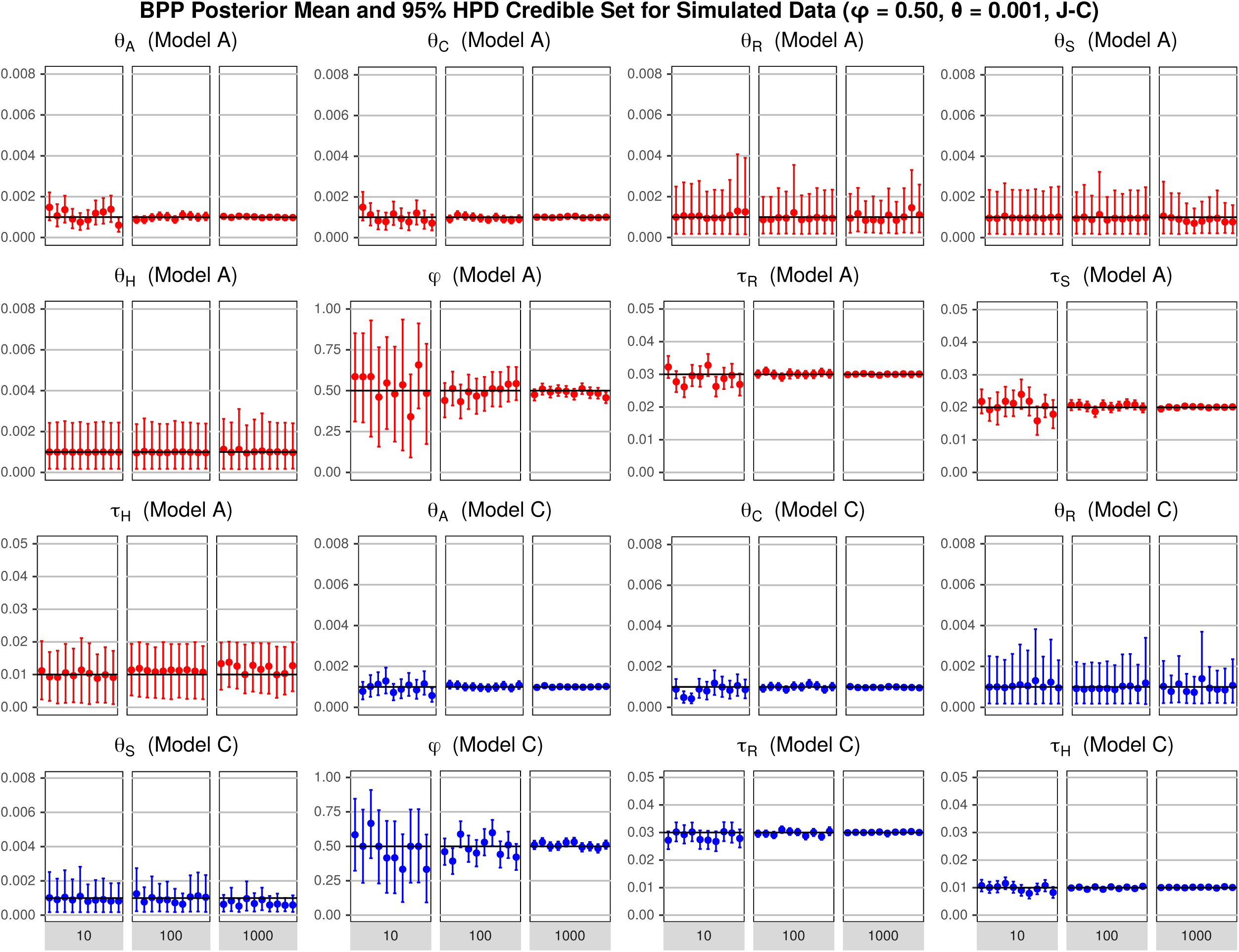
Simulation results for the combination *φ* = 0.5, *θ* = 0.001, and the JC mutation model. See legend to Fig. S1.

**Figure S6:**
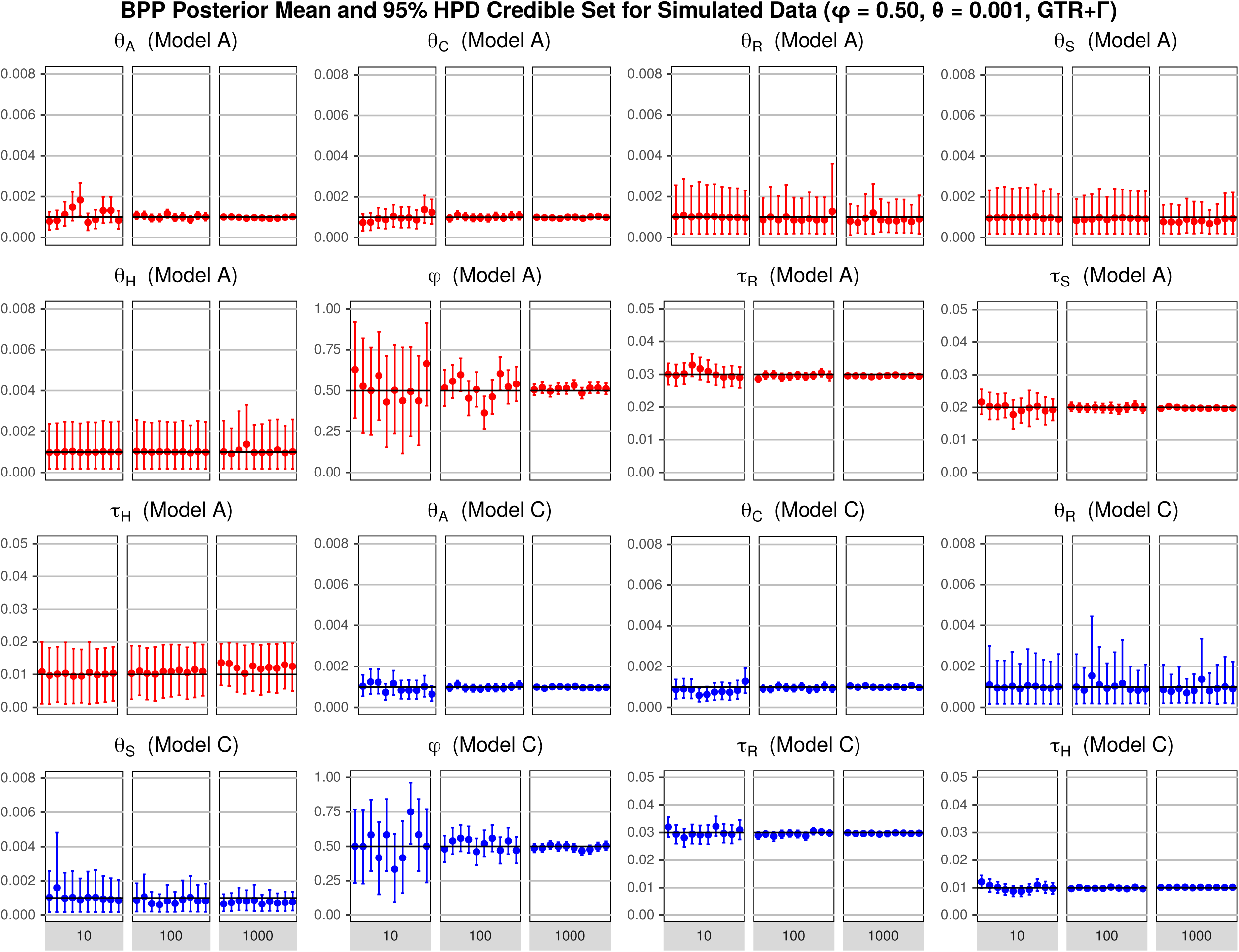
Simulation results for the combination *φ* = 0.5, *θ* = 0.001, and the GTR+Γ mutation model. See legend to Fig. S1.

**Figure S7:**
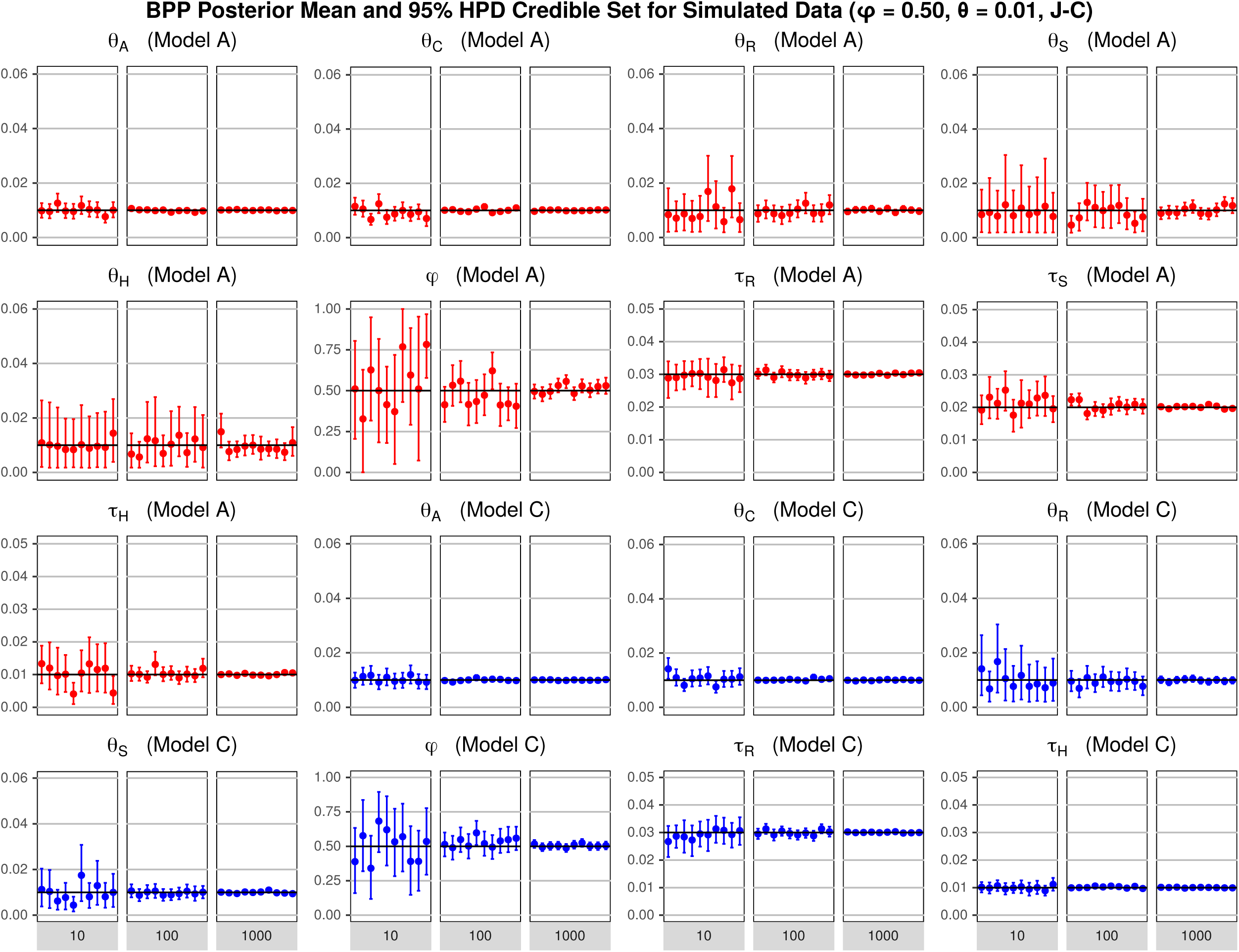
Simulation results for the combination *φ* = 0.5, *θ* = 0.01, and the JC mutation model. See legend to Fig. S1.

**Figure S8:**
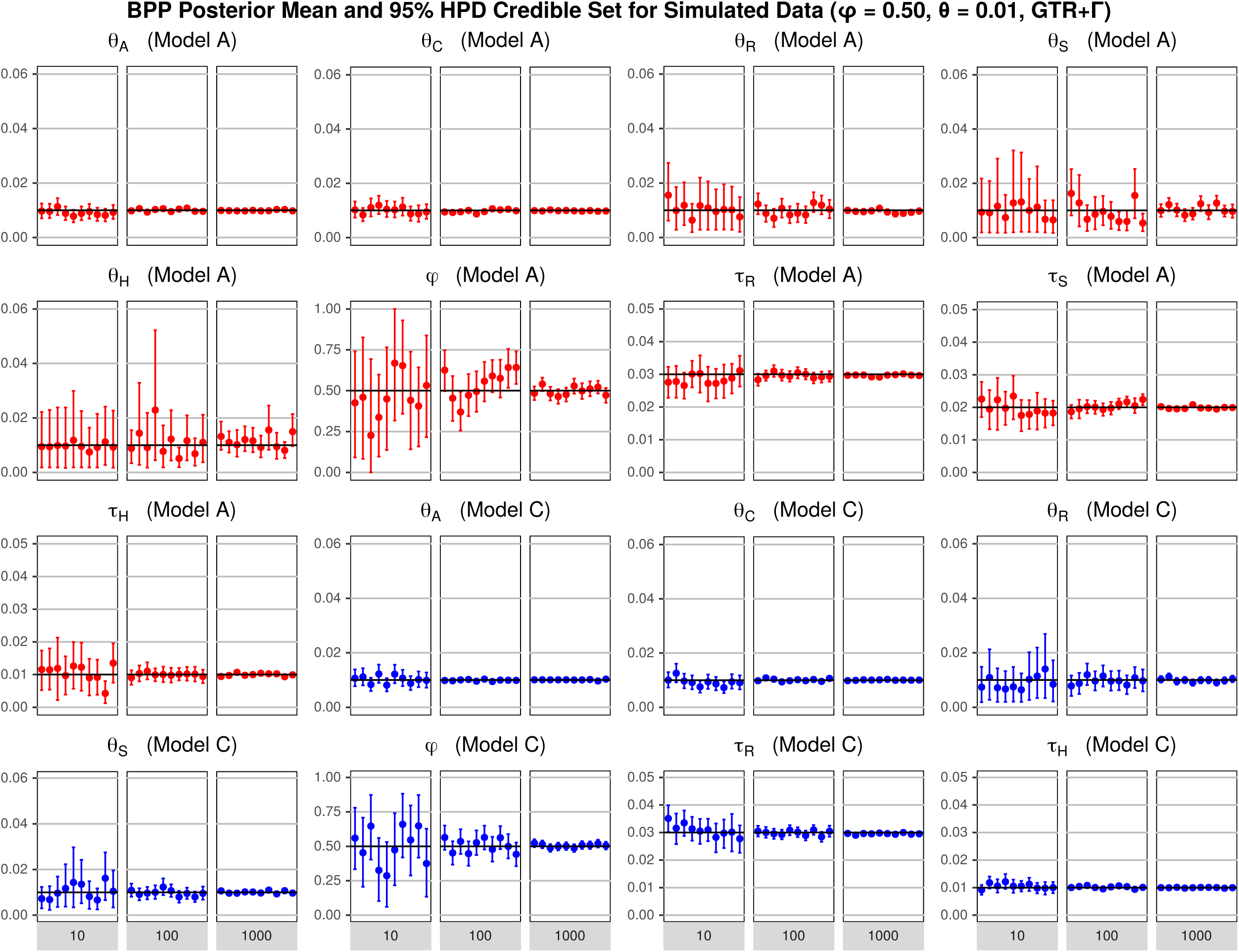
Simulation results for the combination *φ* = 0.5, *θ* = 0.01, and the GTR+Γ mutation model. See legend to Fig. S1.

**Figure S9:**
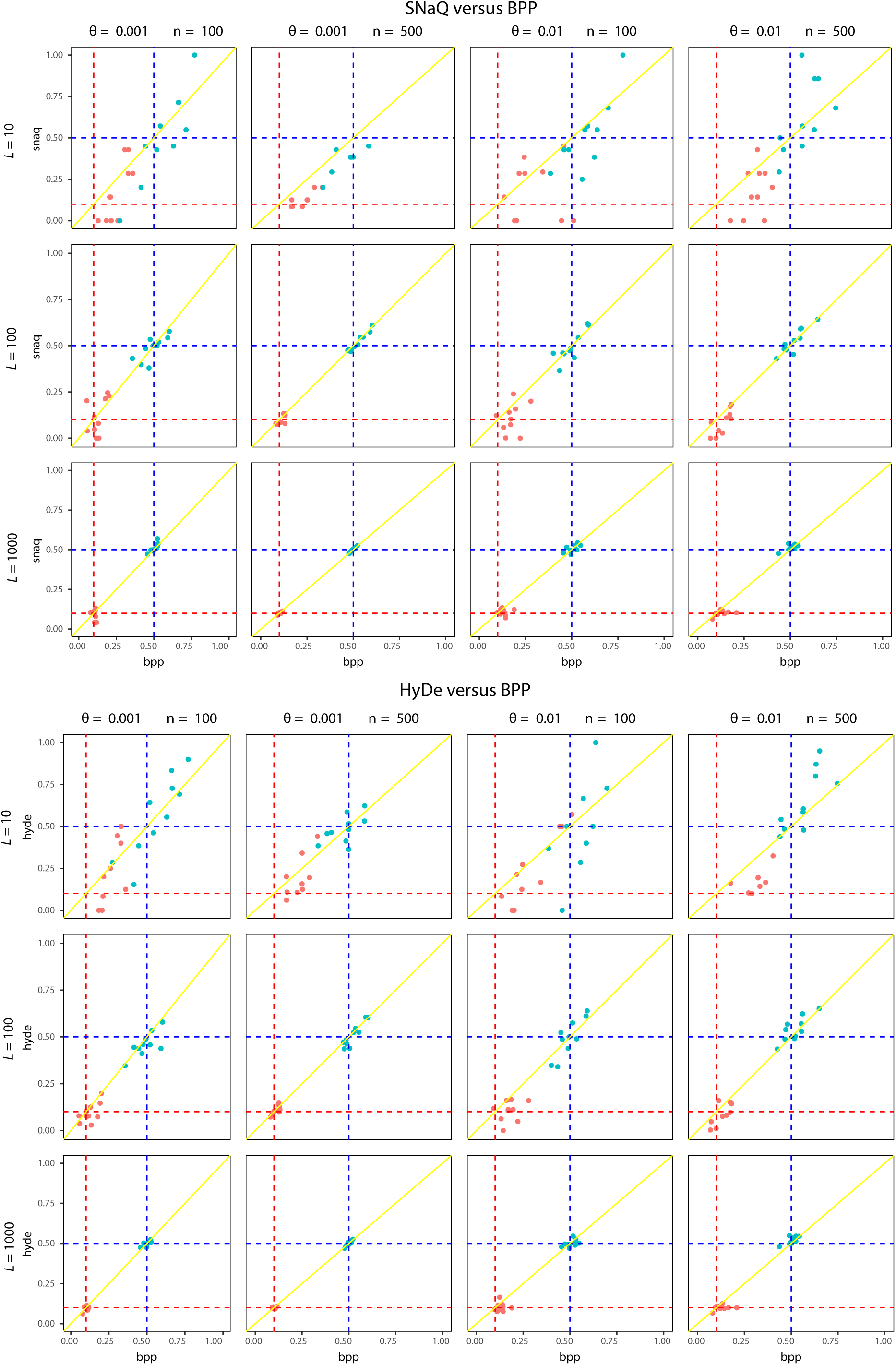
Estimates of *φ* using SNaQ and HyDe plotted against those from BPP on simulated datasets. Model A for three species (Fig. 1A) was used to simulate 10 replicate datasets, using *φ* = 0.1 or 0.5, *θ* = 0.001 or 0.01, the number of loci *L* = 10, 100, or 1000, and sequence length *n* = 100 or 500. The dotted lines indicate the true values.

**Figure S10:**
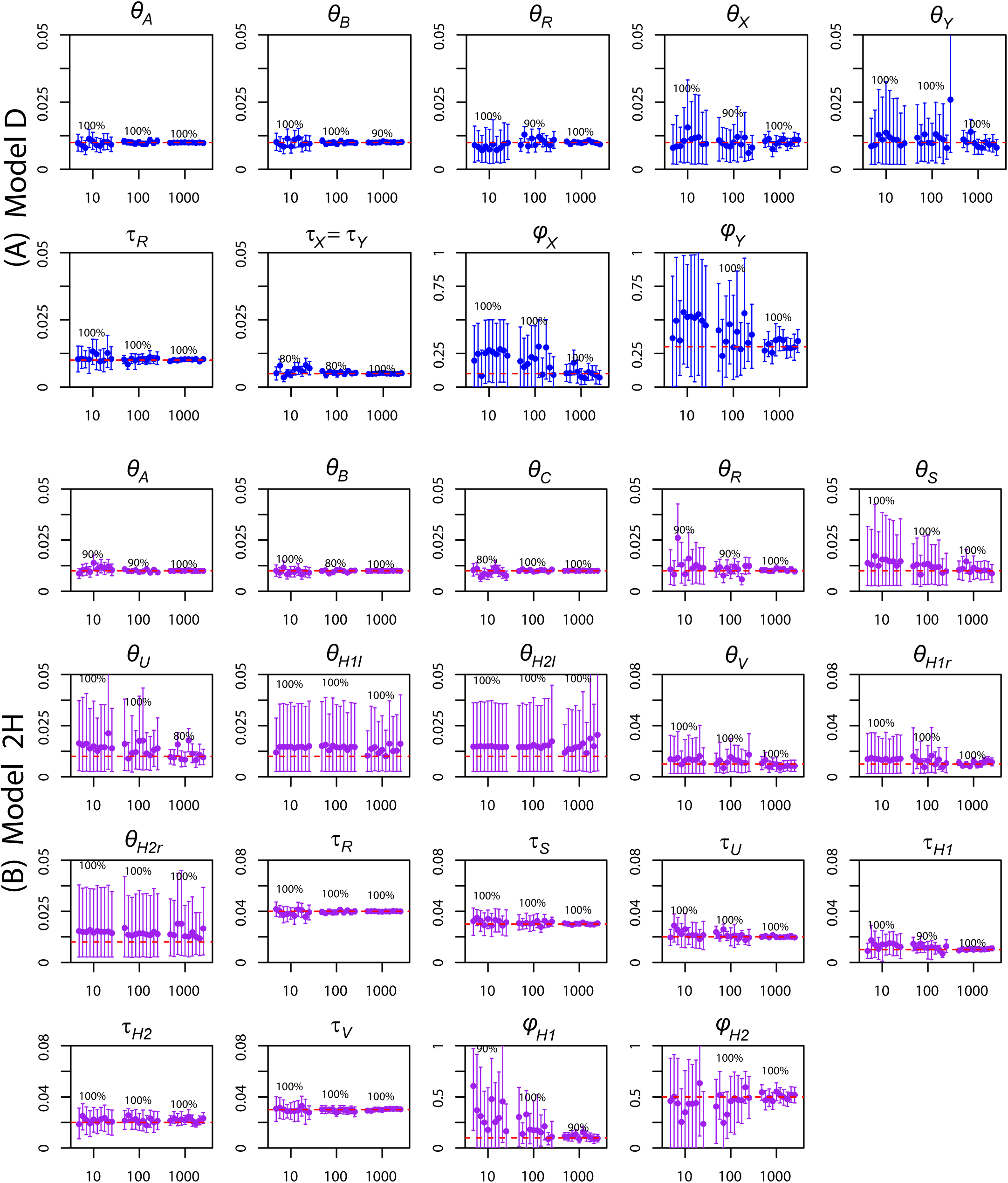
Posterior means and 95% HPD CIs of parameters under model D of Fig. 1 (blue) and model 2H of Fig. 2 (purple) in 10 replicate datasets, each of 10, 100, or 1000 loci. The numbers above the CI bars are the coverage or the proportion of replicate datasets in which the CI bar includes the truth. The horizontal lines represent the true parameter values.

**Table S1.**
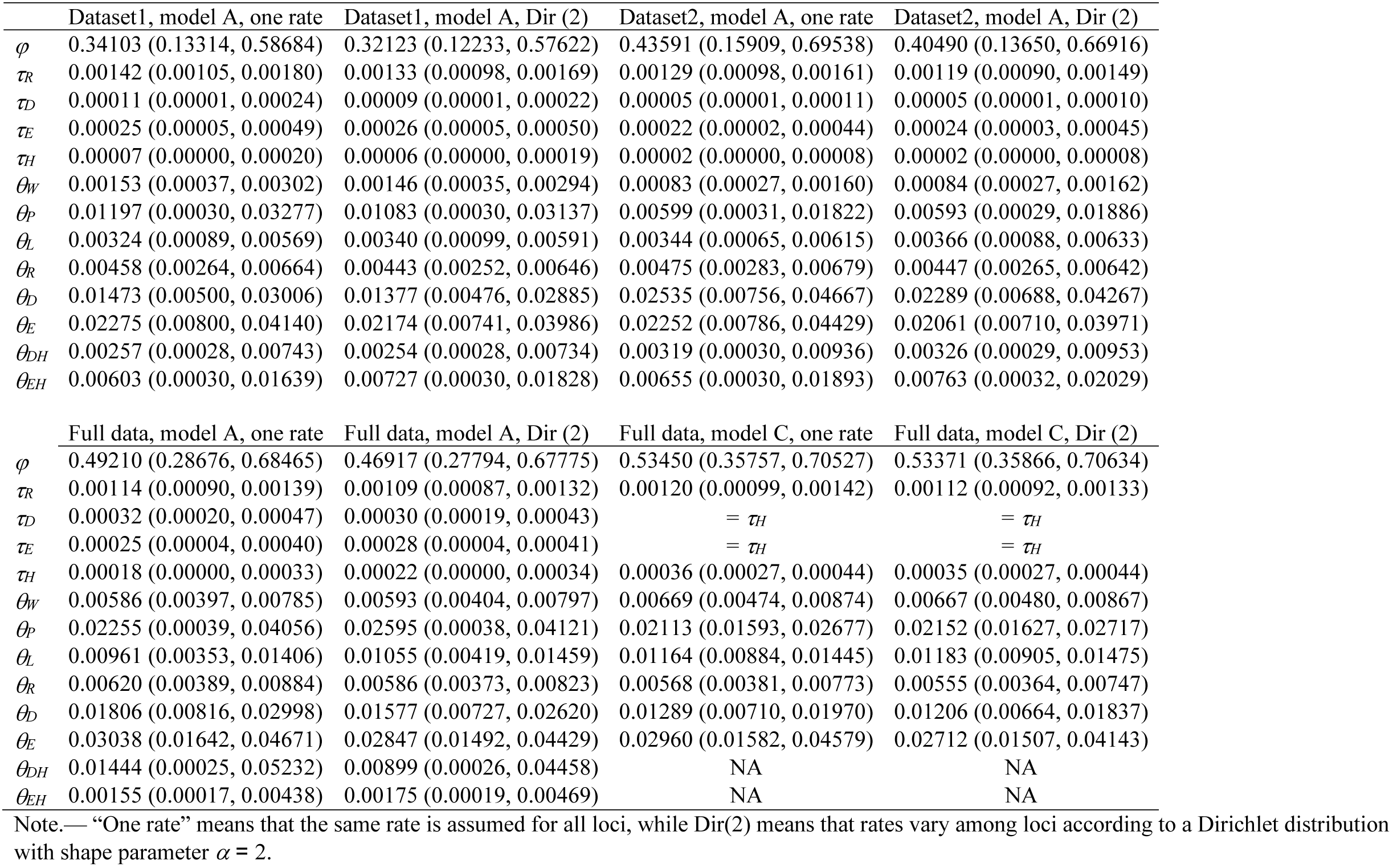
Estimates of parameters under the MSci models A and C of Fig. 1 (posterior means and 95% HPD intervals) for the purple cone spruce datasets

|  | Dataset1, model A, one rate | Dataset1, model A, Dir (2) | Dataset2, model A, one rate | Dataset2, model A, Dir (2) |
| --- | --- | --- | --- | --- |
| $\varphi$ | 0.34103 (0.13314, 0.58684) | 0.32123 (0.12233, 0.57622) | 0.43591 (0.15909, 0.69538) | 0.40490 (0.13650, 0.66916) |
| $\tau_R$ | 0.00142 (0.00105, 0.00180) | 0.00133 (0.00098, 0.00169) | 0.00129 (0.00098, 0.00161) | 0.00119 (0.00090, 0.00149) |
| $\tau_D$ | 0.00011 (0.00001, 0.00024) | 0.00009 (0.00001, 0.00022) | 0.00005 (0.00001, 0.00011) | 0.00005 (0.00001, 0.00010) |
| $\tau_E$ | 0.00025 (0.00005, 0.00049) | 0.00026 (0.00005, 0.00050) | 0.00022 (0.00002, 0.00044) | 0.00024 (0.00003, 0.00045) |
| $\tau_H$ | 0.00007 (0.00000, 0.00020) | 0.00006 (0.00000, 0.00019) | 0.00002 (0.00000, 0.00008) | 0.00002 (0.00000, 0.00008) |
| $\theta_W$ | 0.00153 (0.00037, 0.00302) | 0.00146 (0.00035, 0.00294) | 0.00083 (0.00027, 0.00160) | 0.00084 (0.00027, 0.00162) |
| $\theta_P$ | 0.01197 (0.00030, 0.03277) | 0.01083 (0.00030, 0.03137) | 0.00599 (0.00031, 0.01822) | 0.00593 (0.00029, 0.01886) |
| $\theta_L$ | 0.00324 (0.00089, 0.00569) | 0.00340 (0.00099, 0.00591) | 0.00344 (0.00065, 0.00615) | 0.00366 (0.00088, 0.00633) |
| $\theta_R$ | 0.00458 (0.00264, 0.00664) | 0.00443 (0.00252, 0.00646) | 0.00475 (0.00283, 0.00679) | 0.00447 (0.00265, 0.00642) |
| $\theta_D$ | 0.01473 (0.00500, 0.03006) | 0.01377 (0.00476, 0.02885) | 0.02535 (0.00756, 0.04667) | 0.02289 (0.00688, 0.04267) |
| $\theta_E$ | 0.02275 (0.00800, 0.04140) | 0.02174 (0.00741, 0.03986) | 0.02252 (0.00786, 0.04429) | 0.02061 (0.00710, 0.03971) |
| $\theta_{DH}$ | 0.00257 (0.00028, 0.00743) | 0.00254 (0.00028, 0.00734) | 0.00319 (0.00030, 0.00936) | 0.00326 (0.00029, 0.00953) |
| $\theta_{EH}$ | 0.00603 (0.00030, 0.01639) | 0.00727 (0.00030, 0.01828) | 0.00655 (0.00030, 0.01893) | 0.00763 (0.00032, 0.02029) |
|  | Full data, model A, one rate | Full data, model A, Dir (2) | Full data, model C, one rate | Full data, model C, Dir (2) |
| $\varphi$ | 0.49210 (0.28676, 0.68465) | 0.46917 (0.27794, 0.67775) | 0.53450 (0.35757, 0.70527) | 0.53371 (0.35866, 0.70634) |
| $\tau_R$ | 0.00114 (0.00090, 0.00139) | 0.00109 (0.00087, 0.00132) | 0.00120 (0.00099, 0.00142) | 0.00112 (0.00092, 0.00133) |
| $\tau_D$ | 0.00032 (0.00020, 0.00047) | 0.00030 (0.00019, 0.00043) | $= \tau_H$ | $= \tau_H$ |
| $\tau_E$ | 0.00025 (0.00004, 0.00040) | 0.00028 (0.00004, 0.00041) | $= \tau_H$ | $= \tau_H$ |
| $\tau_H$ | 0.00018 (0.00000, 0.00033) | 0.00022 (0.00000, 0.00034) | 0.00036 (0.00027, 0.00044) | 0.00035 (0.00027, 0.00044) |
| $\theta_W$ | 0.00586 (0.00397, 0.00785) | 0.00593 (0.00404, 0.00797) | 0.00669 (0.00474, 0.00874) | 0.00667 (0.00480, 0.00867) |
| $\theta_P$ | 0.02255 (0.00039, 0.04056) | 0.02595 (0.00038, 0.04121) | 0.02113 (0.01593, 0.02677) | 0.02152 (0.01627, 0.02717) |
| $\theta_L$ | 0.00961 (0.00353, 0.01406) | 0.01055 (0.00419, 0.01459) | 0.01164 (0.00884, 0.01445) | 0.01183 (0.00905, 0.01475) |
| $\theta_R$ | 0.00620 (0.00389, 0.00884) | 0.00586 (0.00373, 0.00823) | 0.00568 (0.00381, 0.00773) | 0.00555 (0.00364, 0.00747) |
| $\theta_D$ | 0.01806 (0.00816, 0.02998) | 0.01577 (0.00727, 0.02620) | 0.01289 (0.00710, 0.01970) | 0.01206 (0.00664, 0.01837) |
| $\theta_E$ | 0.03038 (0.01642, 0.04671) | 0.02847 (0.01492, 0.04429) | 0.02960 (0.01582, 0.04579) | 0.02712 (0.01507, 0.04143) |
| $\theta_{DH}$ | 0.01444 (0.00025, 0.05232) | 0.00899 (0.00026, 0.04458) | NA | NA |
| $\theta_{EH}$ | 0.00155 (0.00017, 0.00438) | 0.00175 (0.00019, 0.00469) | NA | NA |

Note.— “One rate” means that the same rate is assumed for all loci, while Dir(2) means that rates vary among loci according to a Dirichlet distribution with shape parameter $\alpha = 2$ .

**Table S2.** Posterior means (in bold) and 95% HPD intervals (below) of parameters under the MSci model for the Anopheles genomic data (see also table 1)

| | $\theta_{1G}$ | $\theta_{2C}$ | $\theta_{3R}$ | $\theta_{4L}$ | $\theta_{5A}$ | $\theta_{6Q}$ | $\theta_{7o}$ | $\theta_{8g}$ | $\theta_{10a}$ | $\theta_{11c}$ | $\theta_{12d}$ | $\theta_{13e}$ | $\theta_{15b}$ | $\theta_{16h}$ | $\theta_{17f}$ | $\tau_{7o}$ | $\tau_{8g}$ | $\tau_{9h}$ | $\tau_{10a}$ | $\tau_{11c}$ | $\tau_{12d}$ | $\tau_{13e}$ | $\tau_{14f}$ | $\tau_{15b}$ | $\phi_h$ | $\phi_f$ |
| --- | --- | --- | --- | --- | --- | --- | --- | --- | --- | --- | --- | --- | --- | --- | --- | --- | --- | --- | --- | --- | --- | --- | --- | --- | --- | --- |
| **2L1+2 coding** | **4.63** | **2.45** | **0.28** | **0.17** | **0.62** | **0.66** | **0.71** | **2.75** | **46.6** | **0.45** | **0.57** | **0.89** | **1.16** | **2.11** | **3.97** | **0.79** | **0.44** | **0.44** | **0.75** | **0.69** | **0.44** | **0.32** | **0.32** | **0.15** | **0.28** | **0.94** |
|  | 3.39 | 1.96 | 0.27 | 0.16 | 0.59 | 0.61 | 0.66 | 2.00 | 0.36 | 0.29 | 0.49 | 0.77 | 1.01 | 0.32 | 0.77 | 0.76 | 0.40 | 0.40 | 0.70 | 0.67 | 0.40 | 0.31 | 0.31 | 0.13 | 0.24 | 0.92 |
|  | 6.02 | 2.99 | 0.30 | 0.19 | 0.66 | 0.70 | 0.76 | 3.54 | 215.0 | 0.62 | 0.69 | 1.00 | 1.30 | 5.56 | 10.24 | 0.82 | 0.47 | 0.47 | 0.80 | 0.71 | 0.47 | 0.33 | 0.33 | 0.16 | 0.32 | 0.96 |
| **2L1+2 noncoding** | **8.64** | **4.91** | **0.74** | **0.31** | **0.96** | **4.83** | **1.35** | **0.46** | **1.87** | **1.17** | **1.54** | **1.24** | **1.61** | **0.68** | **3.40** | **1.49** | **0.26** | **0.26** | **1.39** | **1.28** | **0.88** | **0.58** | **0.58** | **0.27** | **0.00** | **0.98** |
|  | 7.13 | 4.29 | 0.70 | 0.30 | 0.93 | 3.60 | 1.30 | 0.43 | 0.57 | 0.52 | 1.45 | 1.17 | 1.52 | 0.64 | 0.58 | 1.47 | 0.24 | 0.24 | 1.29 | 1.25 | 0.86 | 0.57 | 0.57 | 0.25 | 0.00 | 0.97 |
|  | 10.28 | 5.56 | 0.79 | 0.32 | 1.00 | 6.21 | 1.39 | 0.49 | 3.17 | 1.63 | 1.64 | 1.31 | 1.71 | 0.72 | 8.20 | 1.51 | 0.29 | 0.29 | 1.49 | 1.30 | 0.89 | 0.59 | 0.59 | 0.28 | 0.00 | 0.99 |
| **2La coding** | **11.41** | **2.36** | **0.73** | **0.24** | **0.81** | **7.67** | **0.85** | **0.24** | **1.53** | **1.98** | **0.28** | **0.53** | **1.97** | **0.47** | **2.12** | **0.76** | **0.18** | **0.18** | **0.60** | **0.60** | **0.58** | **0.24** | **0.24** | **0.19** | **0.01** | **0.73** |
|  | 6.95 | 1.94 | 0.61 | 0.23 | 0.74 | 3.31 | 0.80 | 0.21 | 1.36 | 0.36 | 0.17 | 0.49 | 0.89 | 0.42 | 1.46 | 0.74 | 0.16 | 0.16 | 0.58 | 0.58 | 0.57 | 0.23 | 0.23 | 0.16 | 0.00 | 0.70 |
|  | 16.98 | 2.79 | 0.87 | 0.26 | 0.89 | 13.94 | 0.89 | 0.27 | 1.72 | 4.73 | 0.37 | 0.59 | 3.09 | 0.52 | 2.87 | 0.77 | 0.20 | 0.20 | 0.61 | 0.61 | 0.59 | 0.25 | 0.25 | 0.22 | 0.01 | 0.77 |
| **2La noncoding** | **19.62** | **3.37** | **1.10** | **0.41** | **1.22** | **8.91** | **1.14** | **0.46** | **2.22** | **3.03** | **2.11** | **0.73** | **1.49** | **0.65** | **2.91** | **1.69** | **0.38** | **0.38** | **1.14** | **1.14** | **1.14** | **0.42** | **0.42** | **0.39** | **0.00** | **0.64** |
|  | 15.53 | 3.11 | 1.03 | 0.39 | 1.16 | 6.17 | 1.10 | 0.43 | 2.12 | 0.39 | 0.37 | 0.70 | 0.30 | 0.61 | 2.56 | 1.67 | 0.35 | 0.35 | 1.13 | 1.13 | 1.13 | 0.41 | 0.41 | 0.37 | 0.00 | 0.63 |
|  | 24.02 | 3.65 | 1.18 | 0.41 | 1.28 | 12.2 | 1.18 | 0.50 | 2.32 | 7.70 | 5.22 | 0.77 | 2.73 | 0.70 | 3.27 | 1.71 | 0.41 | 0.41 | 1.16 | 1.16 | 1.15 | 0.43 | 0.43 | 0.41 | 0.00 | 0.65 |
| **2R coding** | **6.75** | **1.87** | **0.25** | **0.16** | **0.74** | **0.66** | **0.61** | **3.30** | **2.07** | **0.28** | **0.58** | **0.85** | **0.95** | **1.80** | **4.71** | **0.70** | **0.35** | **0.35** | **0.67** | **0.64** | **0.35** | **0.28** | **0.28** | **0.17** | **0.34** | **0.97** |
|  | 4.98 | 1.63 | 0.24 | 0.15 | 0.66 | 0.61 | 0.57 | 2.55 | 0.53 | 0.18 | 0.51 | 0.06 | 0.16 | 0.36 | 0.74 | 0.68 | 0.31 | 0.31 | 0.66 | 0.63 | 0.31 | 0.23 | 0.23 | 0.14 | 0.30 | 0.96 |
|  | 8.78 | 2.13 | 0.27 | 0.17 | 0.81 | 0.71 | 0.64 | 4.05 | 5.14 | 0.38 | 0.64 | 1.79 | 1.41 | 4.28 | 12.87 | 0.74 | 0.38 | 0.38 | 0.69 | 0.65 | 0.38 | 0.33 | 0.33 | 0.22 | 0.39 | 0.98 |
| **2R noncoding** | **9.85** | **3.32** | **0.59** | **0.33** | **1.25** | **1.37** | **0.66** | **7.51** | **1.24** | **1.61** | **0.92** | **1.57** | **1.83** | **2.07** | **12.75** | **1.84** | **0.90** | **0.90** | **1.36** | **1.32** | **0.90** | **0.53** | **0.53** | **0.30** | **0.22** | **0.97** |
|  | 8.69 | 3.13 | 0.55 | 0.32 | 1.21 | 1.25 | 0.07 | 2.82 | 0.88 | 0.20 | 0.80 | 1.44 | 1.71 | 0.35 | 1.56 | 1.52 | 0.75 | 0.75 | 1.26 | 1.26 | 0.75 | 0.52 | 0.52 | 0.29 | 0.12 | 0.96 |
|  | 11.06 | 3.53 | 0.63 | 0.34 | 1.29 | 1.48 | 1.36 | 14.4 | 1.49 | 5.15 | 1.15 | 1.75 | 1.95 | 5.17 | 38.38 | 2.11 | 1.00 | 1.00 | 1.63 | 1.38 | 1.00 | 0.55 | 0.55 | 0.30 | 0.34 | 0.99 |
| **3L1+2 coding** | **3.02** | **1.71** | **0.30** | **0.19** | **0.56** | **0.50** | **0.78** | **1.38** | **1.01** | **1.38** | **0.41** | **0.91** | **0.88** | **2.48** | **6.40** | **0.72** | **0.43** | **0.43** | **0.69** | **0.67** | **0.43** | **0.22** | **0.22** | **0.06** | **0.32** | **0.94** |
|  | 1.34 | 1.02 | 0.27 | 0.17 | 0.51 | 0.46 | 0.72 | 0.97 | 0.35 | 0.28 | 0.34 | 0.80 | 0.75 | 0.34 | 1.17 | 0.69 | 0.39 | 0.39 | 0.66 | 0.65 | 0.39 | 0.20 | 0.20 | 0.05 | 0.27 | 0.92 |
|  | 5.35 | 2.53 | 0.32 | 0.20 | 0.61 | 0.54 | 0.84 | 1.83 | 2.06 | 3.57 | 0.48 | 1.02 | 1.00 | 5.53 | 15.92 | 0.75 | 0.45 | 0.45 | 0.72 | 0.70 | 0.45 | 0.23 | 0.23 | 0.07 | 0.37 | 0.96 |
| **3L1+2 noncoding** | **7.63** | **2.70** | **0.58** | **0.36** | **0.95** | **0.89** | **1.51** | **5.64** | **1.58** | **0.61** | **0.69** | **1.60** | **1.67** | **2.10** | **8.11** | **1.37** | **0.84** | **0.84** | **1.32** | **1.28** | **0.84** | **0.39** | **0.39** | **0.13** | **0.33** | **0.96** |
|  | 4.90 | 2.21 | 0.56 | 0.34 | 0.90 | 0.86 | 1.45 | 4.32 | 0.53 | 0.33 | 0.63 | 1.51 | 1.54 | 0.33 | 1.90 | 1.35 | 0.82 | 0.82 | 1.28 | 1.25 | 0.82 | 0.38 | 0.38 | 0.11 | 0.30 | 0.95 |
|  | 10.99 | 3.23 | 0.60 | 0.37 | 1.00 | 0.93 | 1.57 | 7.07 | 3.18 | 0.89 | 0.75 | 1.69 | 1.80 | 5.30 | 17.93 | 1.40 | 0.86 | 0.86 | 1.37 | 1.30 | 0.87 | 0.40 | 0.40 | 0.14 | 0.36 | 0.97 |
| **3La coding** | **14.96** | **6.68** | **0.40** | **0.20** | **1.07** | **1.16** | **0.84** | **0.78** | **1.66** | **1.73** | **0.53** | **1.64** | **1.71** | **1.99** | **5.92** | **0.68** | **0.38** | **0.38** | **0.67** | **0.67** | **0.38** | **0.21** | **0.21** | **0.13** | **0.65** | **0.93** |
|  | 4.95 | 3.43 | 0.38 | 0.19 | 0.95 | 1.06 | 0.79 | 0.67 | 0.32 | 0.32 | 0.46 | 1.39 | 0.98 | 0.37 | 1.26 | 0.66 | 0.36 | 0.36 | 0.64 | 0.63 | 0.36 | 0.20 | 0.20 | 0.10 | 0.62 | 0.91 |
|  | 31.55 | 11.03 | 0.41 | 0.22 | 1.20 | 1.26 | 0.88 | 0.89 | 4.06 | 4.23 | 0.59 | 1.91 | 2.49 | 4.92 | 14.15 | 0.70 | 0.39 | 0.39 | 0.70 | 0.70 | 0.39 | 0.23 | 0.23 | 0.16 | 0.68 | 0.95 |
| **3La noncoding** | **189.3** | **16.95** | **0.94** | **0.38** | **1.56** | **1.91** | **0.48** | **1.54** | **2.64** | **1.88** | **0.86** | **2.07** | **3.87** | **2.13** | **3.73** | **2.05** | **0.93** | **0.93** | **1.46** | **1.45** | **0.93** | **0.46** | **0.46** | **0.24** | **0.54** | **0.98** |
|  | 39.12 | 11.80 | 0.81 | 0.35 | 1.47 | 1.67 | 0.01 | 0.98 | 0.44 | 0.29 | 0.69 | 1.71 | 3.08 | 0.37 | 0.73 | 1.61 | 0.65 | 0.65 | 1.37 | 1.37 | 0.65 | 0.43 | 0.43 | 0.22 | 0.46 | 0.97 |
|  | 634.5 | 22.84 | 1.03 | 0.42 | 1.66 | 2.07 | 1.44 | 2.49 | 5.23 | 5.27 | 1.12 | 2.74 | 4.59 | 5.25 | 9.00 | 2.27 | 1.10 | 1.10 | 1.61 | 1.60 | 1.10 | 0.48 | 0.48 | 0.27 | 0.70 | 0.98 |
| **3R coding** | **8.32** | **4.14** | **0.31** | **0.20** | **0.84** | **0.81** | **0.76** | **1.97** | **2.92** | **0.37** | **0.85** | **4.78** | **0.69** | **1.98** | **6.40** | **0.74** | **0.32** | **0.32** | **0.71** | **0.67** | **0.32** | **0.25** | **0.25** | **0.17** | **0.43** | **0.95** |
|  | 5.54 | 3.22 | 0.28 | 0.17 | 0.75 | 0.71 | 0.72 | 1.48 | 0.83 | 0.25 | 0.68 | 1.11 | 0.17 | 0.34 | 1.17 | 0.72 | 0.26 | 0.26 | 0.68 | 0.66 | 0.26 | 0.21 | 0.21 | 0.13 | 0.37 | 0.93 |
|  | 11.66 | 5.13 | 0.33 | 0.22 | 0.93 | 0.91 | 0.80 | 2.44 | 6.60 | 0.48 | 1.01 | 12.91 | 1.31 | 4.90 | 16.89 | 0.77 | 0.39 | 0.39 | 0.73 | 0.68 | 0.39 | 0.29 | 0.29 | 0.21 | 0.49 | 0.96 |
| **3R noncoding** | **15.27** | **6.68** | **0.87** | **0.36** | **1.48** | **4.08** | **0.92** | **0.56** | **1.99** | **0.39** | **1.75** | **1.59** | **2.20** | **0.55** | **6.12** | **1.61** | **0.51** | **0.51** | **1.31** | **1.30** | **0.82** | **0.56** | **0.56** | **0.25** | **0.03** | **0.98** |
|  | 12.35 | 5.96 | 0.79 | 0.35 | 1.43 | 2.20 | 0.06 | 0.46 | 1.09 | 0.10 | 1.63 | 1.51 | 2.09 | 0.21 | 0.76 | 1.38 | 0.38 | 0.38 | 1.27 | 1.25 | 0.81 | 0.55 | 0.55 | 0.24 | 0.01 | 0.97 |
|  | 18.49 | 7.43 | 0.93 | 0.38 | 1.52 | 5.56 | 1.41 | 0.71 | 3.01 | 0.70 | 1.90 | 1.68 | 2.31 | 0.75 | 16.46 | 2.02 | 0.73 | 0.73 | 1.38 | 1.37 | 0.83 | 0.57 | 0.57 | 0.26 | 0.06 | 0.98 |

Note.— Estimates of $\theta$ s and $\tau$ s are multiplied by 100.

## References

Arnold, B. J., Lahner, B., DaCosta, J. M., Weisman, C. M., Hollister, J. D., Salt, D. E., Bomblies, K., and Yant, L. 2016. Borrowed alleles and convergence in serpentine adaptation. Proc. Natl. Acad. SciU.S.A., 113(29): 8320–8325.

Blischak, P. D., Chifman, J., Wolfe, A. D., and Kubatko, L. S. 2018. Hyde: A python package for genome-scale hybridization detection. Syst. Biol., 67(5): 821–829.

Burgess, R. and Yang, Z. 2008. Estimation of hominoid ancestral population sizes under Bayesian coalescent models incorporating mutation rate variation and sequencing errors. Mol. Biol. Evol., 25(9): 1979–1994.

Cardona, G., Rossello, F., and Valiente, G. 2008. Extended Newick: it is time for a standard representation of phylogenetic networks. BMC Bioinformatics, 9: 532.

Chan, Y. C., Roos, C., Inoue-Murayama, M., Inoue, E., Shih, C. C., Pei, K. J., and Vigilant, L. 2013. Inferring the evolutionary histories of divergences in *Hylobates* and *Nomascus* gibbons through multilocus sequence data. BMC Evol. Biol., 13: 82.

Dalquen, D., Zhu, T., and Yang, Z. 2017. Maximum likelihood implementation of an isolation-with-migration model for three species. Syst. Biol., 66: 379–398.

Degnan, J. H. 2018. Modeling hybridization under the network multispecies coalescent. Syst. Biol., 67(5): 786–799.

Durand, E. Y., Patterson, N., Reich, D., and Slatkin, M. 2011. Testing for ancient admixture between closely related populations. Mol. Biol. Evol., 28: 2239–2252.

Edwards, S. V., Xi, Z., Janke, A., Faircloth, B. C., McCormack, J. E., Glenn, T. C., Zhong, B., Wu, S., Lemmon, E. M., Lemmon, A. R., Leaché, A. D., Liu, L., and Davis, C. C. 2016. Implementing and testing the multispecies coalescent model: A valuable paradigm for phylogenomics. Mol. Phylogenet. Evol., 94(Pt A): 447–462.

Ellegren, H., Smeds, L., Burri, R., Olason, P. I., Backstrom, N., Kawakami, T., Kunstner, A., Makinen, H., Nadachowska-Brzyska, K., Qvarnstrom, A., Uebbing, S., and Wolf, J. B. W. 2012. The genomic landscape of species divergence in *Ficedula* flycatchers. Nature, 491: 756–760.

Felsenstein, J. 1981. Evolutionary trees from DNA sequences: a maximum likelihood approach. J. Mol. Evol., 17(6): 368–376.

Flouris, T., Jiao, X., Rannala, B., and Yang, Z. 2018. Species tree inference with BPP using genomic sequences and the multispecies coalescent. Mol. Biol. Evol., 35(10): 2585–2593.

Folk, R. A., Soltis, P. S., Soltis, D. E., and Guralnick, R. 2018. New prospects in the detection and comparative analysis of hybridization in the tree of life. Am. J. Bot., 105(3): 364–375.

Fontaine, M. C., Pease, J. B., Steele, A., Waterhouse, R. M., Neafsey, D. E., Sharakhov, I. V., Jiang, X., Hall, A. B., Catteruccia, F., Kakani, E., Mitchell, S. N., Wu, Y. C., Smith, H. A., Love, R. R., Lawniczak, M. K., Slotman, M. A., Emrich, S. J., Hahn, M. W., and Besansky, N. J. 2015. Extensive introgression in a malaria vector species complex revealed by phylogenomics. Science, 347(6217): 1258524.

Green, R. E., Krause, J., Briggs, A. W., Maricic, T., Stenzel, U., Kircher, M., Patterson, N., Li, H., Zhai, W., Fritz, M. H., Hansen, N. F., Durand, E. Y., Malaspinas, A. S., Jensen, J. D., Marques-Bonet, T., Alkan, C., Prufer, K., Meyer, M., Burbano, H. A., Good, J. M., Schultz, R., Aximu-Petri, A., Butthof, A., Hober, B., Hoffner, B., Siegemund, M., Weihmann, A., Nusbaum, C., Lander, E. S., Russ, C., Novod, N., Affourtit, J., Egholm, M., Verna, C., Rudan, P., Brajkovic, D., Kucan, Z., Gusic, I., Doronichev, V. B., Golovanova, L. V., Lalueza-Fox, C., de la Rasilla, M., Fortea, J., Rosas, A., Schmitz, R. W., Johnson, P. L., Eichler, E. E., Falush, D., Birney, E., Mullikin, J. C., Slatkin, M., Nielsen, R., Kelso, J., Lachmann, M., Reich, D., and Paabo, S. 2010. A draft sequence of the neandertal genome. Science, 328: 710–722.

Harrison, R. G. and Larson, E. L. 2014. Hybridization, introgression, and the nature of species boundaries. J. Hered., 105 (S1): 795–809.

Hey, J. 2010. Isolation with migration models for more than two populations. Mol. Biol. Evol., 27: 905–920.

Hey, J. and Nielsen, R. 2004. Multilocus methods for estimating population sizes, migration rates and divergence time, with applications to the divergence of *Drosophila pseudoobscura* and *D. persimilis*. Genetics, 167: 747–760.

Hey, J., Chung, Y., Sethuraman, A., Lachance, J., Tishkoff, S., Sousa, V. C., and Wang, Y. 2018. Phylogeny estimation by integration over isolation with migration models. Mol. Biol. Evol., 35(11): 2805–2818.

Huson, D. H., Rupp, R., and Cornavacca, C. 2011. Phylogenetic Networks: Concepts, Algorithms and Applications. Cambridge University Press, Cambridge, England.

Jackson, N. D., Carstens, B. C., Morales, A. E., and O’Meara, B. C. 2017. Species delimitation with gene flow. Syst. Biol., 66(5): 799–812.

Jones, G. R. 2019. Divergence estimation in the presence of incomplete lineage sorting and migration. Syst. Biol., 68(1): 19–31.

Jukes, T. and Cantor, C. 1969. Evolution of protein molecules. In H. Munro, editor, Mammalian Protein Metabolism, pages 21–123. Academic Press, New York.

Kubatko, L. S. 2009. Identifying hybridization events in the presence of coalescence via model selection. Syst. Biol., 58(5): 478–488.

Kumar, V., Lammers, F., Bidon, T., Pfenninger, M., Kolter, L., Nilsson, M. A., and Janke, A. 2017. The evolutionary history of bears is characterized by gene flow across species. Sci Rep, 7: 46487.

Lartillot, N. and Philippe, H. 2006. Computing Bayes factors using thermodynamic integration. Syst. Biol., 55: 195–207.

Leaché, A. D., Zhu, T., Rannala, B., and Yang, Z. 2019. The spectre of too many species. Syst. Biol., 68(1): 168–181.

Liu, S., Lorenzen, E. D., Fumagalli, M., Li, B., Harris, K., Xiong, Z., Zhou, L., Korneliussen, T. S., Somel, M., Babbitt, C., Wray, G., Li, J., He, W., Wang, Z., Fu, W., Xiang, X., Morgan, C. C., Doherty, A., O’Connell, M. J., McInerney, J. O., Born, E. W., Dalen, L., Dietz, R., Orlando, L., Sonne, C., Zhang, G., Nielsen, R., Willerslev, E., and Wang, J. 2014. Population genomics reveal recent speciation and rapid evolutionary adaptation in polar bears. Cell, 157: 785–794.

Lohse, K., Chmelik, M., Martin, S. H., and Barton, N. H. 2016. Efficient strategies for calculating blockwise likelihoods under the coalescent. Genetics, 202(2): 775–786.

Mallet, J., Besansky, N., and Hahn, M. W. 2016. How reticulated are species? BioEssays, 38(2): 140–149.

Mao, Y., Economo, E. P., and Satoh, N. 2018. The roles of introgression and climate change in the rise to dominance of *Acropora* corals. Curr. Biol., 28(21): 3373–3382 e5.

Martin, S. H. and Jiggins, C. D. 2017. Interpreting the genomic landscape of introgression. Curr. Opin. Genet. Dev., 47: 69–74.

Martin, S. H., Dasmahapatra, K. K., Nadeau, N. J., Salazar, C., Walters, J. R., Simpson, F., Blaxter, M., Manica, A., Mallet, J., and Jiggins, C. D. 2013. Genome-wide evidence for speciation with gene flow in *Heliconius* butterflies. Genome Res., 23(11): 1817–1828.

Nielsen, R., Akey, J. M., Jakobsson, M., Pritchard, J. K., Tishkoff, S., and Willerslev, E. 2017. Tracing the peopling of the world through genomics. Nature, 541: 302.

O’Hagan, A. and Forster, J. 2004. Kendall’s Advanced Theory of Statistics: Bayesian Inference. Arnold, London.

Rannala, B. and Yang, Z. 2003. Bayes estimation of species divergence times and ancestral population sizes using DNA sequences from multiple loci. Genetics, 164(4): 1645–1656.

Rannala, B. and Yang, Z. 2017. Efficient bayesian species tree inference under the multispecies coalescent. Syst. Biol., 66: 823–842.

Shi, C. and Yang, Z. 2018. Coalescent-based analyses of genomic sequence data provide a robust resolution of phylogenetic relationships among major groups of gibbons. Mol. Biol. Evol., 35: 159–179.

Slotman, M. A., Della Torre, A., Calzetta, M., and Powell, J. R. 2005. Differential introgression of chromosomal regions between *Anopheles gambiae* and *An. arabiensis*. Am. J. Trop. Med. Hyg., 73(2): 326–335.

Solis-Lemus, C. and Ane, C. 2016. Inferring phylogenetic networks with maximum pseudolikelihood under incomplete lineage sorting. PLoS Genet., 12(3): e1005896.

Solis-Lemus, C., Bastide, P., and Ane, C. 2017. Phylonetworks: A package for phylogenetic networks. Mol. Biol. Evol., 34(12): 3292–3298.

Stamatakis, A., Aberer, A., Goll, C., Smith, S., Berger, S., and Izquierdo-Carrasco, F. 2012. Raxml-light: a tool for computing terabyte phylogenies. Bioinformatics, 28: 2064–2066.

Sun, Y., Abbott, R. J., Li, L., Li, L., Zou, J., and Liu, J. 2014. Evolutionary history of purple cone spruce (picea purpurea) in the qinghai-tibet plateau: homoploid hybrid origin and pleistocene expansion. Mol. Ecol., 23(2): 343–359.

Thawornwattana, Y., Dalquen, D., and Yang, Z. 2018a. Coalescent analysis of phylogenomic data confidently resolves the species relationships in the *Anopheles gambiae* species complex. Mol. Biol. Evol., 35(10): 2512–2527.

Thawornwattana, Y., Dalquen, D., and Yang, Z. 2018b. Designing simple and efficient Markov chain Monte Carlo proposal kernels. Bayesian Analysis, 13(4): 1033–1059.

Wen, D. and Nakhleh, L. 2018. Coestimating reticulate phylogenies and gene trees from multilocus sequence data. Syst. Biol., 67(3): 439–457.

Wu, D.-D., Ding, X.-D., Wang, S., Wojcik, J. M., Zhang, Y.,Tokarska, M., Li, Y., Wang, M.-S., Faruque, O., Nielsen, R., Zhang, Q., and Zhang, Y.-P. 2018. Pervasive introgression facilitated domestication and adaptation in the bos species complex. Nature Ecol. Evol., 2(7): 1139–1145.

Xu, B. and Yang, Z. 2016. Challenges in species tree estimation under the multispecies coalescent model. Genetics, 204: 1353–1368. doi:10.1534/genetics.116.190173.

Yang, Z. 1994a. Estimating the pattern of nucleotide substitution. J. Mol. Evol., 39(1): 105–11.

Yang, Z. 1994b. Maximum likelihood phylogenetic estimation from DNA sequences with variable rates over sites: approximate methods. J. Mol. Evol., 39: 306–314.

Yang, Z. 2015. The BPP program for species tree estimation and species delimitation. Curr. Zool., 61: 854–865.

Yang, Z. and Rodriguez, C. E. 2013. Searching for efficient Markov chain Monte Carlo proposal kernels. Proc. Natl. Acad. Sci. U.S.A., 110(48): 19307–19312.

Yu, Y., Dong, J., Liu, K. J., and Nakhleh, L. 2014. Maximum likelihood inference of reticulate evolutionary histories. Proc. Natl. Acad. Sci. U.S.A., 111(46): 16448–16453.

Zhang, C., Ogilvie, H. A., Drummond, A. J., and Stadler, T. 2018. Bayesian inference of species networks from multilocus sequence data. Mol. Biol. Evol., 35: 504–517.

Zhu, S. and Degnan, J. H. 2017. Displayed trees do not determine distinguishability under the network multispecies coalescent. Syst. Biol., 66(2): 283–298.

Zhu, T. and Yang, Z. 2012. Maximum likelihood implementation of an isolation-with-migration model with three species for testing speciation with gene flow. Mol. Biol. Evol., 29: 3131–3142.

